# Metatranscriptomics reveals diverse microbial communities in PCR-negative severe acute respiratory infections

**DOI:** 10.64898/2026.03.19.712874

**Authors:** Nicola S Holdsworth, Rebecca K French, Stephanie J Waller, Lauren Jelley, Meaghan P O’Neill, Isa de Vries, Jérémy Dubrulle, Nigel French, Max Bloomfield, David Winter, Q. Sue Huang, Jemma L Geoghegan

**Affiliations:** Department of Microbiology and Immunology, Faculty of Biomedical Sciences, University of Otago, Dunedin, 9016, New Zealand; New Zealand Institute for Public Health and Forensic Science, Wellington, New Zealand; Tāwharau Ora/School of Veterinary Science, Massey University, Palmerston North, New Zealand; Department of Microbiology, Awanui Laboratories, Wellington, New Zealand

**Keywords:** SARI, ARI, Virome, Metagenomics, Infectome, Undiagnosed infection, New Zealand

## Abstract

Severe acute respiratory infections (SARI) are a major cause of hospitalisation and mortality globally, yet a proportion of cases remain without an identified aetiology following routine diagnostic testing. Here, we used metatranscriptomic sequencing to characterise microbial communities in 300 PCR-negative SARI nasopharyngeal samples collected through sentinel hospital-based surveillance in Aotearoa New Zealand between 2014 and 2021. Transcripts from microbial species with recognised or potential associations with respiratory infection were detected in 43% of samples, comprising 10 RNA viruses, three DNA viruses, nine bacterial species and four fungal species. Multiple microbial species were detected in 26% of positive samples, highlighting the polymicrobial composition of a subset of PCR-negative SARI cases. Human rhinoviruses were the most frequently detected viruses despite being included in the original PCR screening, while seasonal human coronaviruses, human parechovirus A1 and parainfluenza virus type 4 were also identified, including viruses not included in the diagnostic panel. Bacterial and fungal taxa associated with respiratory infection were also detected, alongside transcripts corresponding to characterised virulence factors and antimicrobial resistance genes. Infectome composition showed statistically significant but modest associations with patient age and year of sample collection. As healthy comparative samples were unavailable, detection of microbial RNA cannot establish causality or distinguish infection from colonisation or incidental detection. Nevertheless, these findings demonstrate the additional microbial and genomic information obtainable through unbiased sequencing of diagnostically unresolved respiratory samples and support further evaluation of metatranscriptomics as a complementary approach to targeted PCR for respiratory disease surveillance and investigation.

## Introduction

Accurate and timely diagnosis of respiratory infections remains a major challenge in clinical medicine (1–4). Despite widespread adoption of PCR-based diagnostic panels (5–11), a substantial proportion of patients hospitalised with severe acute respiratory infections (SARI) receive no identifiable aetiology. These targeted assays are highly sensitive for known pathogens and are widely implemented in diagnostic laboratories using high-throughput, kit-based platforms. The pathogens included in these panels typically represent the most common and clinically relevant causes of disease. However, because panel composition varies between instruments and laboratories, and is necessarily restricted to predefined targets, such assays cannot detect organisms not included in the diagnostic kits, nor reliably identify rare, unexpected, or novel microbes (12). In addition, they provide limited insight into co-infections or the broader microbial communities that may influence disease severity.

This diagnostic gap is clinically significant in Aotearoa New Zealand, where rates of respiratory hospitalisation are high and disproportionately affect Māori and Pacific peoples (13,14). The national SARI surveillance system, established through the Southern Hemisphere Influenza and Vaccine Effectiveness Research and Surveillance (SHIVERS) programme, relies on PCR and serology to characterise circulating respiratory pathogens (15). While effective for routine surveillance of common viruses such as influenza viruses, respiratory syncytial virus (RSV), severe acute respiratory syndrome coronavirus 2 (SARS-CoV-2), enterovirus, adenovirus, human metapneumovirus (hMPV), parainfluenza viruses 1-3 and rhinoviruses, these methods are pathogen specific and often leave the cause of some SARI cases unresolved, limiting the information available for clinical decision-making.

High-throughput metagenomic sequencing offers an opportunity to overcome these constraints (16). Specifically, metatranscriptomic sequencing provides an unbiased approach that captures all actively transcribing organisms in a clinical sample, enabling simultaneous detection of viral, bacterial and fungal species, including those not targeted by routine diagnostics (17). By also revealing functional information such as potential virulence factors and antimicrobial resistance gene expression, metatranscriptomics can greatly improve understanding of complex or polymicrobial infections (18).

Here, we apply metatranscriptomic sequencing to PCR-negative SARI cases collected through the SHIVERS surveillance programme to (i) characterise the microbial communities present in diagnostically unresolved SARI samples in New Zealand; (ii) identify viral, bacterial and fungal taxa not captured by routine viral testing; (iii) investigate demographic and temporal factors associated with infectome composition; and (iv) characterise antimicrobial resistance and virulence-associated gene signatures. This study provides the first metatranscriptomic characterisation of PCR-negative SARI samples in New Zealand and evaluates the additional information that unbiased sequencing can provide alongside targeted respiratory surveillance.

## Methods

### Ethics statement

The NZ Northern A Health and Disability Ethics Committee approved the SHIVERS studies (NTX/11/11/102). No formal consent from individuals was obtained because the data are anonymous.

### Sample collection

SHIVERS hospital-based SARI surveillance was located within two District Health Boards of the Auckland region of New Zealand: Auckland District Health Board (ADHB) and Counties Manukau District Health Board (CMDHB). This is a predominantly urban population of ∼906,000 people, with a wide spectrum of socio-economic, cultural, ethnic and demographic groups broadly representative of the New Zealand population. Between 2014 and 2021, 1,274 patient samples were collected from ADHB and CMDHB hospitals. Patients who met the World Health Organisation’s (WHO) case definition for severe acute respiratory infection (SARI) (19) (i.e. an acute respiratory infection with a history of fever or measured fever of ≥38 °C and cough with onset within the last 10 days, requiring hospitalisation) were enrolled in the study.

Respiratory specimens, mainly nasopharyngeal swabs, were obtained and referred to the New Zealand Institute for Public Health and Forensic Science (PHF Science) for testing. Viral RNA was extracted either manually using the Zymo ZR Viral RNA kit (R1035) or using the Thermofisher™ Magmax Viral/Pathogen nucleic acid isolation kit (A48383) as per manufacturer’s instructions on the ZiXpress or Kingfisher Flex extraction platforms. Samples underwent respiratory viral testing by real-time reverse transcription PCR (rRT-PCR) using the AgPath-ID™ One-Step RT-PCR kit (4387424). Singleplex primers and probes were used to test for influenza A and B viruses (20), adenovirus, RSV, hMPV, parainfluenza viruses 1–3, human rhinovirus and enterovirus (21) (Figure 1). SARS-CoV-2 viral testing was added to the testing algorithm from 2020 onwards using the CCDC primers and probes for N gene coupled with the Quantbio qscript XLT 1 step-RT-qPCR ToughMix™ (95132–500). Of the 1,274 samples that tested negative for all viral targets as listed above, 300 were selected for metatranscriptomic analysis based on total RNA concentration of >1.5 ng/µl.

**Figure 1.**
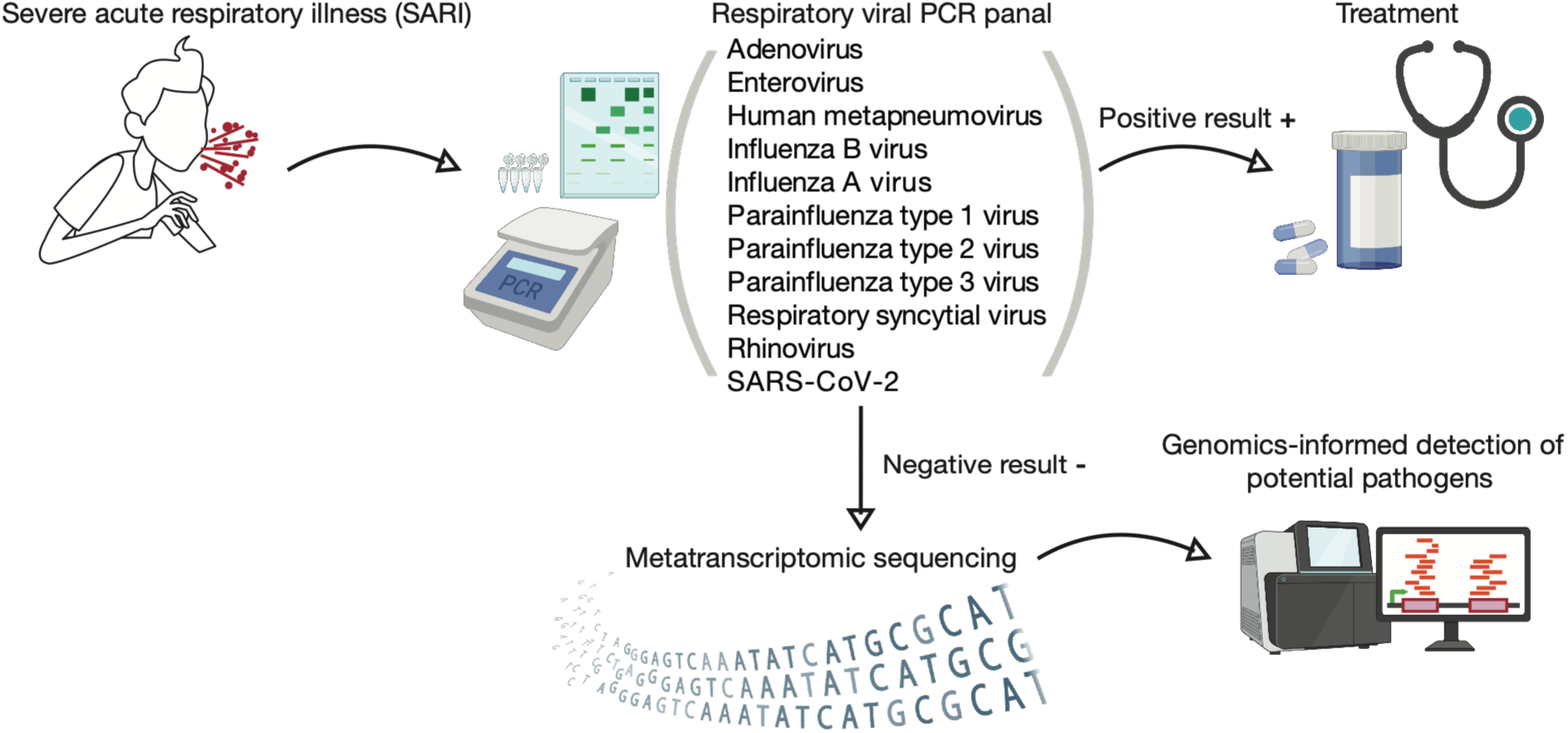
Workflow for microbial identification in SARI patient samples. Nasopharyngeal samples taken from patients meeting the SARI criteria were first tested using a PCR viral panel. Samples negative by PCR were selected for metatranscriptomic sequencing, enabling detection and characterisation of viral, bacterial and fungal species beyond the predefined targets of the initial PCR panel.

### Total RNA extraction and sequencing

Total RNA was re-extracted from 300 patient samples using the ZymoBIOMICS MagBead RNA kit (Zymo Research), strictly following the manufacturer’s protocol without any modifications. RNA concentration was quantified using a NanoDrop spectrophotometer, with an average concentration of 5.5 ng/µl, ranging from 1.5 ng/µl to 78 ng/µl. Each sample was subject to library preparation using the Illumina Stranded Total RNA Prep with Ribo-Zero Plus kit (Illumina), which depletes both cytoplasmic and mitochondrial ribosomal RNA. Amplification of the libraries was performed with 16 PCR cycles. Paired-end (150 bp) sequencing was conducted on the Illumina NovaSeqX platform.

### Quality control, assembly and annotation

Low quality bases were trimmed using a sliding window approach, and Nextera paired-end adapters were removed using Trimmomatic (v0.38) (22). Following trimming, reads were assessed for quality using FastQC (v0.11.8) (23). Reads aligning to the human genome (GRCh38/hg38) using Bowtie2 (v2.4.4) (24) were removed and excluded from downstream analysis. The remaining reads were assembled de novo into contigs using MEGAHIT (v1.2.9) (25).

### Viral identification and abundance estimation

Assembled contigs were subject to sequence similarity searches against a local copy of the NCBI non-redundant protein database using DIAMOND BLASTx (v2.1.9) (26). To reduce false positives, we used a maximum expected e-value of 1 × 10^−10^ to filter for putative viral contigs. The BLASTx ‘sskingdoms’ output field was used to assist assignment of contigs to broad taxonomic kingdoms. During viral screening, all contigs assigned to non-viral taxa were manually removed. Putative viral contigs were further examined at the nucleotide level using Geneious Prime (v2025.0.3).

Viral abundance was quantified by mapping reads to the assembled viral contigs using Bowtie2 (24). To normalise for variation in sequencing depth across libraries, abundance was expressed as reads per million (RPM), calculated by dividing the number of viral reads by the total number of reads in the corresponding library and multiplying by one million. Potential cross sample contamination due to index hopping was identified if viral reads representing less than 0.1% of the total read count in a library shared greater than 99% nucleotide identity with sequences in another library (27). No viral sequences meeting these criteria were identified.

### Virus phylogenetic analysis

Nucleotide sequences of putative viral transcripts were aligned using the automatic algorithm selection in MAFFT (v7.490) (28) with their closest genetic relatives identified using BLASTn, as well as representative nucleotide sequences from the same taxonomic species retrieved from GenBank. Rhinovirus species were further classified by genotype using an automated web-based enterovirus typing tool based on the VP1 gene (29). IQ-TREE (v1.6.12) (30) was used to estimate maximum likelihood phylogenetic trees for each viral species using ModelFinder (31) with 1,000 ultra-fast bootstrapping replicates. Phylogenetic trees were annotated using Figtree (v1.2.2) (32). SnapGene (v8.2) (33) was used to align the rhinovirus primer sequence used by PHF Science during initial PCR testing against the identified rhinovirus sequences to assess congruence and coverage.

### Exploring non-viral microorganisms

CCMetagen (v1.4.3) (34) was used to classify non-viral sequence reads and calculate abundances (in RPM) at the species level. A curated version of the NCBI nucleotide database prepared by CCMetagen was downloaded and pre-indexed using k-mer alignment (KMA) (v1.4.15) (35) required by CCMetagen. Reads were first mapped to this database using KMA and alignment data were processed by CCMetagen to generate taxonomic abundance profiles for each library.

### Statistical analysis

Associations between the presence of viral, bacterial, and fungal organisms and demographic variables (age group and sex) were assessed using Fisher’s exact test (36) or the chi-squared test (37), depending on the data. Fisher’s exact test was used when organism prevalence was low or when contingency tables contained sparse cell counts. The chi-squared test was applied only to comparisons where all expected counts in the contingency table were sufficiently large, such as analysis restricted to the 129 samples with at least one microbial species detected. All tests were two-sided, and statistical significance was evaluated using a threshold of p < 0.05.

### Alpha diversity of infectome composition

Alpha diversity within each sample was assessed using abundance (standardised RPM), richness (number of microbial species), and the Shannon and Simpson diversity indices. Abundance, Shannon and Simpson indices were analysed using the Kruskal-Wallis rank sum test (38) to assess differences across groups. Where significant, post-hoc pairwise comparisons were performed using Dunn’s test with Bonferroni adjustment (39), which adjusts p-values when multiple tests are performed on the same dataset to minimise the rate of false positives (40). A generalised linear model with a Poisson distribution was used to test for differences in richness.

### Beta diversity of infectome composition

Beta diversity was assessed at the microbial species level using the phyloseq package in R (41). A permutational multivariate analysis of variance (PERMANOVA) via the adonis2 function in the vegan package (42) was used to test for statistical significance (p < 0.05) of the effect of patient age, location, sex and year of sample collection on infectome beta diversity. The results were visualised using principal coordinates analysis (PCoA) ordination plots based on the Bray-Curtis matrix, measuring the dissimilarity of microbial communities between libraries ranging from 0 (identical communities) to 1 (no microbial species in common).

### Uncovering bacterial virulence factors

Virulence factors for the four most abundant bacterial species were identified by aligning trimmed reads to annotated complete reference genomes using Bowtie2 and quantifying abundance with the Rsubread (v2.14.2) R package (43). We assessed the expression of *Moraxella catarrhalis*, *Serratia marcescens*, *Staphylococcus aureus* and *Streptococcus pneumoniae* genes using the following reference sequences: GCA_002080125.1, GCA_030291735.1, GCA_000013425.1 and GCA_001457635.1, respectively. Known virulence genes were curated from the Virulence Factors of Pathogenic Bacteria (VFDB) database (44). As this approach is reference-dependent, only virulence factors present in the selected reference genomes were evaluated. Consequently, strain-specific or accessory virulence genes not represented in these references may not have been captured.

### Assessing the resistome

Trimmed reads were aligned to the ResFinder database using KMA (k-mer alignment) (45) to identify antimicrobial resistance genes (ARG). This analysis was run for all identified bacterial species against the entire ResFinder database of ARGs.

## Results

### Undiagnosed SARI patients

A total of 300 samples were selected (see Methods) from patients presenting with SARI between May 2014 and June 2021 whose molecular diagnosis remained elusive following a pan-viral PCR-based panel comprising tests for Influenza (A and B) virus, adenovirus, RSV, hMPV, parainfluenza viruses 1-3, human rhinovirus, SARS-CoV-2 and enterovirus. Of these patients, 151 (50%) were female, 146 (49%) were male and 3 (1%) had an unspecified sex (Figure 2). A total of 201 samples were obtained from ADHB hospital and 99 from CMDHB hospital. Patient age was well distributed across the range, including 64 patients (21%) under 5 years of age, 111 (37%) between 5 and 65 years, and 102 (34%) over 65 years old. Age information was not available for 23 patients (8%) (Figure 2). It is important to note that the temporal distribution of samples was uneven. A large proportion of samples were collected in 2015, whereas fewer samples were available during the COVID-19 pandemic (from 2020). Therefore, the 300 samples should be interpreted as a subset of PCR-negative SARI cases over this period and may not fully represent the total SARI burden in New Zealand.

**Figure 2.**
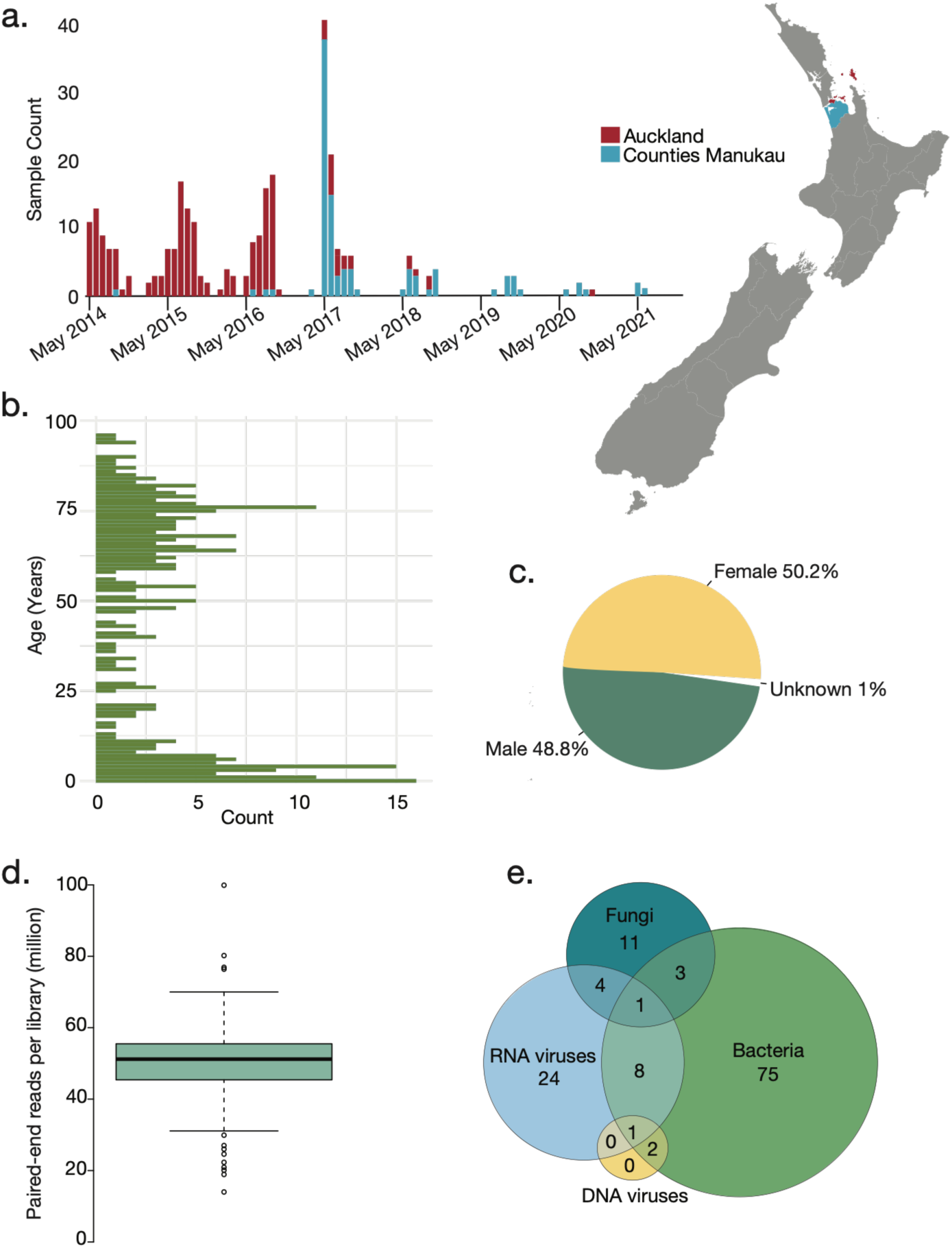
SARI patient meta data and high-level results. (a) Bar graph illustrating sample count over time and map of New Zealand highlighting the regions of SARI patient sample collection. (b) Histogram showing the distribution of patient age. (c) Pie chart showing sex distribution among patients. (d) Box plot showing the distribution of sequencing reads (millions) among sequencing libraries, showing the median and interquartile range. (e) Venn diagram representing the number of viral, bacterial and fungal sequences found in SARI patients, including samples containing multiple microbial species.

### The nasopharyngeal metatranscriptome of undiagnosed SARI patients

RNA sequencing of total RNA from nasopharyngeal swabs collected from 300 SARI patients in New Zealand generated an average of 50.2 million raw reads per library, ranging from 14 million to 100 million (Figure 2). One sample (17HB0134) was excluded from further analysis due to low sequence coverage (2.8 million raw reads). The overall metatranscriptome across the remaining 299 samples consisted primarily of bacterial sequences (∼80%), followed by eukaryotes (∼19%), viral sequences (∼0.15%) and archaea (∼0.0004%).

### The total infectome

A metatranscriptomic analysis of nasopharyngeal samples successfully identified a range of viral, bacterial, and fungal transcripts. For this study, we focused on pathogens associated with human respiratory disease, including species with established pathogenicity. An abundance threshold of 1 RPM was applied to define microbial presence. Although the analysis allowed for the potential detection of novel pathogens, no previously unreported species were identified in this cohort.

All microbes identified and characterised in this study had been previously described in humans, with the analysis focused on taxa with recognised or potential associations with respiratory infection. In total, we identified 10 RNA viruses, three DNA viruses, nine bacterial species and four fungal species (Supplementary Table 1). Out of 299 patient samples, 129 (43%) contained at least one microbial species meeting our inclusion criteria (Figure 2e). Of these, 38 patient samples contained one or more RNA viruses, three contained one or more DNA viruses, 90 contained one or more bacterial species, and 19 samples had one or more fungal species. Multiple microbial species were detected in 26% (33 of 129) of positive samples and could comprise multiple species of the same type (e.g. two bacteria) or different microbial types (bacteria, viruses or fungi). Co-detection involving at least two different microbial types was observed in 19 cases: seven in children aged 0-5 years, five in individuals aged 5-65 years, three in those aged 65 and older and four in patients with unknown age (Supplementary Table 2). The most frequently detected species across these samples were *Streptococcus pneumoniae* (identified in 34 samples), *Pseudomonas aeruginosa* (22 samples) and *Staphylococcus aureus* (20 samples). In terms of total abundance across all samples, the most predominant species identified in this study were *Serratia marcescens*, *Human coronavirus OC43*, *Naganishia albida* and *Purpureocillium lilacinum* (Figure 3).

**Figure 3.**
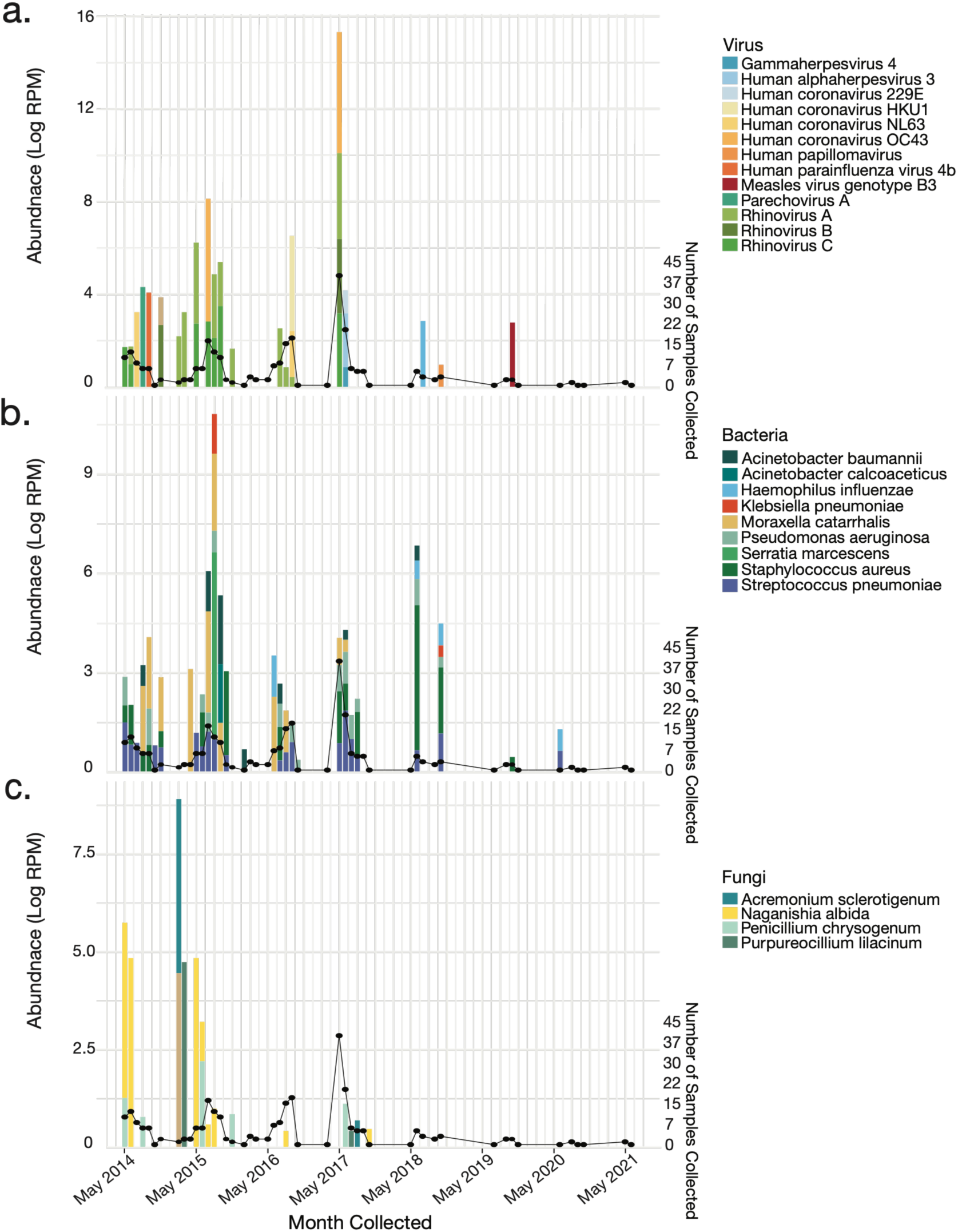
Total infectome abundance. Stacked log abundance of identified viral (a), bacterial (b), and fungal (c) species across all patient samples, with black dotted line representing number of patient samples collected each month. Month and year collected shown on the bottom of panel (c) for clarity but corresponds to all panels.

### Viruses associated with SARI

Virus detection overall (n = 299) varied significantly by age group (Fisher’s exact test, p = 0.0002). When expressed as the proportion within each age group, children aged 0-5 years had the highest proportion of virus-positive samples (29.8%), followed by 5-65 years (8.9%) and adults >65 years (7.2%). Detection did not differ significantly by sex (Fisher’s exact test, p = 0.095). In total, we detected 13 virus species across five viral families (Supplementary table 3), with the most common being human rhinoviruses A-C, found in 23 individuals (Figure 4). Of these, 13 were classified as Human rhinovirus A, with nine further assigned to genogroups A8, A18, A20, A22, A34, A46, A68, A81 and A101. We identified Human rhinovirus B in one individual, classified as genogroup B3, and a further nine individuals with Human rhinovirus C, falling across genogroups C1, C2, C3, C15, C26, C32, C36 and C56. Overall, we recovered six full genomes, while the others were partial sequences. Among the individuals with human rhinoviruses, 46% were aged 0-5 years, 29% were aged 5-65 years, while 12.5% were aged >65 and 12.5% were of an unknown age. We detected the greatest number of rhinoviruses in 2015 (n = 12) and 2017 (n = 5), where together these years accounted for 74% of identifications. Among the 13 cases of rhinovirus A identified, co-infection was detected in five (39%), which consisted of three different fungi and two different bacteria species. Of the nine cases of rhinovirus C, five (55%) occurred as co-infections, three of which were co-infected with Moraxella catarrhalis. The single case of rhinovirus B was a co-infection with three species (*Streptococcus pneumoniae*, *Moraxella catarrhalis* and *Human papillomavirus*). Co-infection was most common in patients aged 0-5 years, with four of ten cases (40%) affected. The average abundance of the 13 rhinovirus A cases was 857 RPM, compared with 756 RPM for the nine rhinovirus C cases. The most abundant rhinovirus species was Human rhinovirus A (11,152 RPM), which overall was the fifth most abundant viral species detected in this study. This was followed by Human rhinovirus C (7,569 RPM), and Human rhinovirus B (471 RPM). SnapGene was used to compare the primer sequence used by PHF Science against our longer rhinovirus sequences (>1,000 bp), where consistent complementarity was observed across all sequences examined, including Human rhinovirus A (n = 3), Human rhinovirus B (n = 1), and Human rhinovirus C (n = 6) (Supplementary Figure 1).

**Figure 4.**
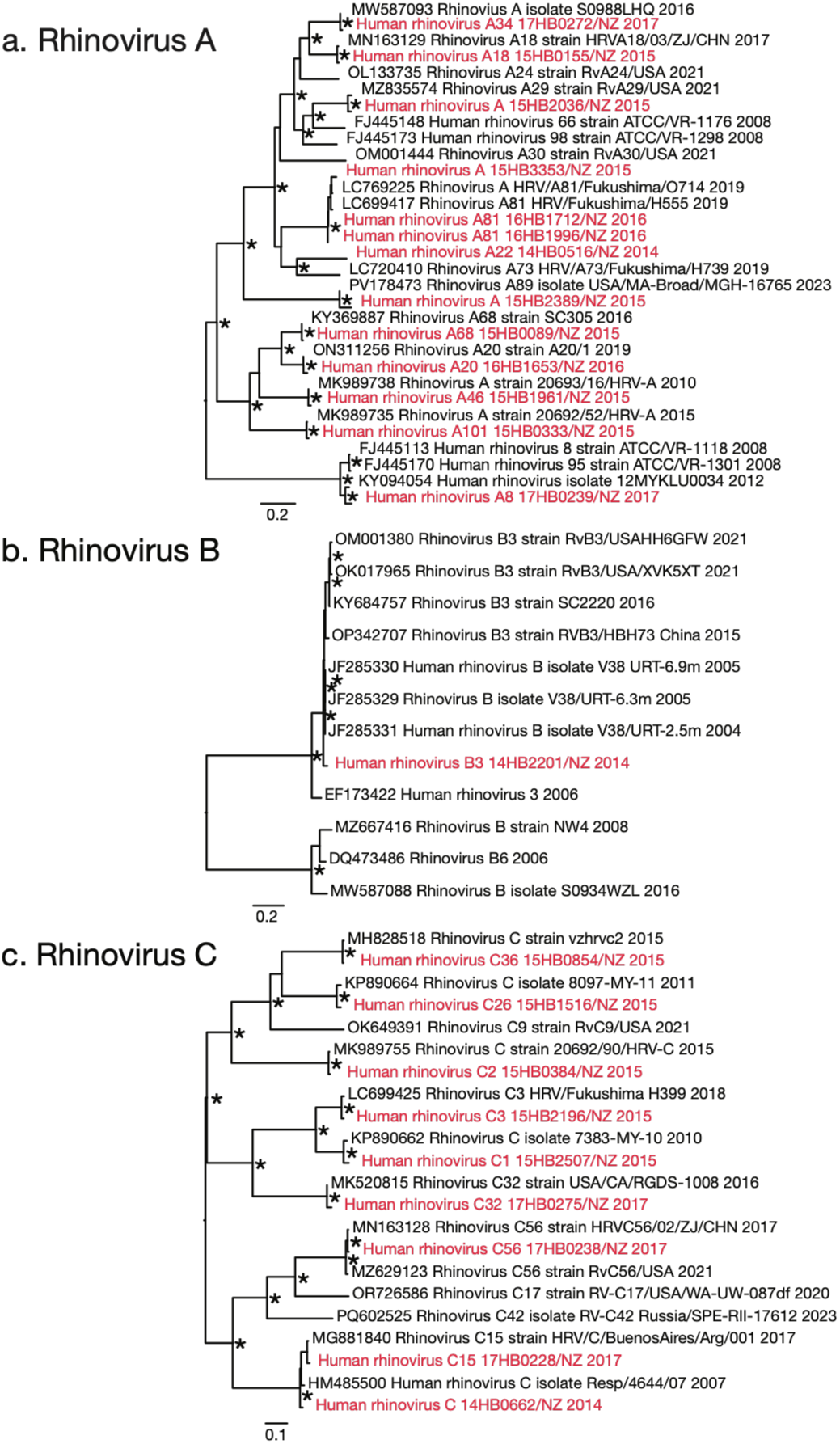
Maximum likelihood phylogenetic trees of human rhinovirus species. Maximum likelihood phylogenetic trees of human rhinoviruses identified (red) and representative nucleotide sequences obtained from NCBI taxonomy (black). Sequences identified in this study are marked in red representing, (a) Human rhinovirus A, (b) Human rhinovirus B and (c) Human rhinovirus C. Phylogenetic trees are midpoint rooted for clarity, and nodes with bootstrap values of >95% are noted with asterisks. Branches are scaled by number of nucleotide substitutions per site shown in the scale bar.

All four types of seasonal human coronavirus (hCoV) were present in this dataset. We found Human coronavirus 229E (n = 2), Human coronavirus OC43 (n = 5), Human coronavirus HKU1 (n = 2) and Human coronavirus NL63 (n = 2) (Figure 5). Among these sequences, the overall abundances ranged from 9-169,164 RPM, and we identified four complete genomes. Of the individuals with a coronavirus detected, 36% were children (0-5 years), 27% were aged between 5-65, while 27% were aged >65, and 9% were of an unknown age. Human coronaviruses were detected across multiple years (2014–2018), with the highest number of detections recorded in 2015 (n = 5). Of the 11 coronavirus-positive samples, 82% (n = 9) were sampled in Auckland and 18% (n = 2) in Counties Manukau. Additional bacterial species were detected in two samples containing Human coronavirus OC43.

**Figure 5.**
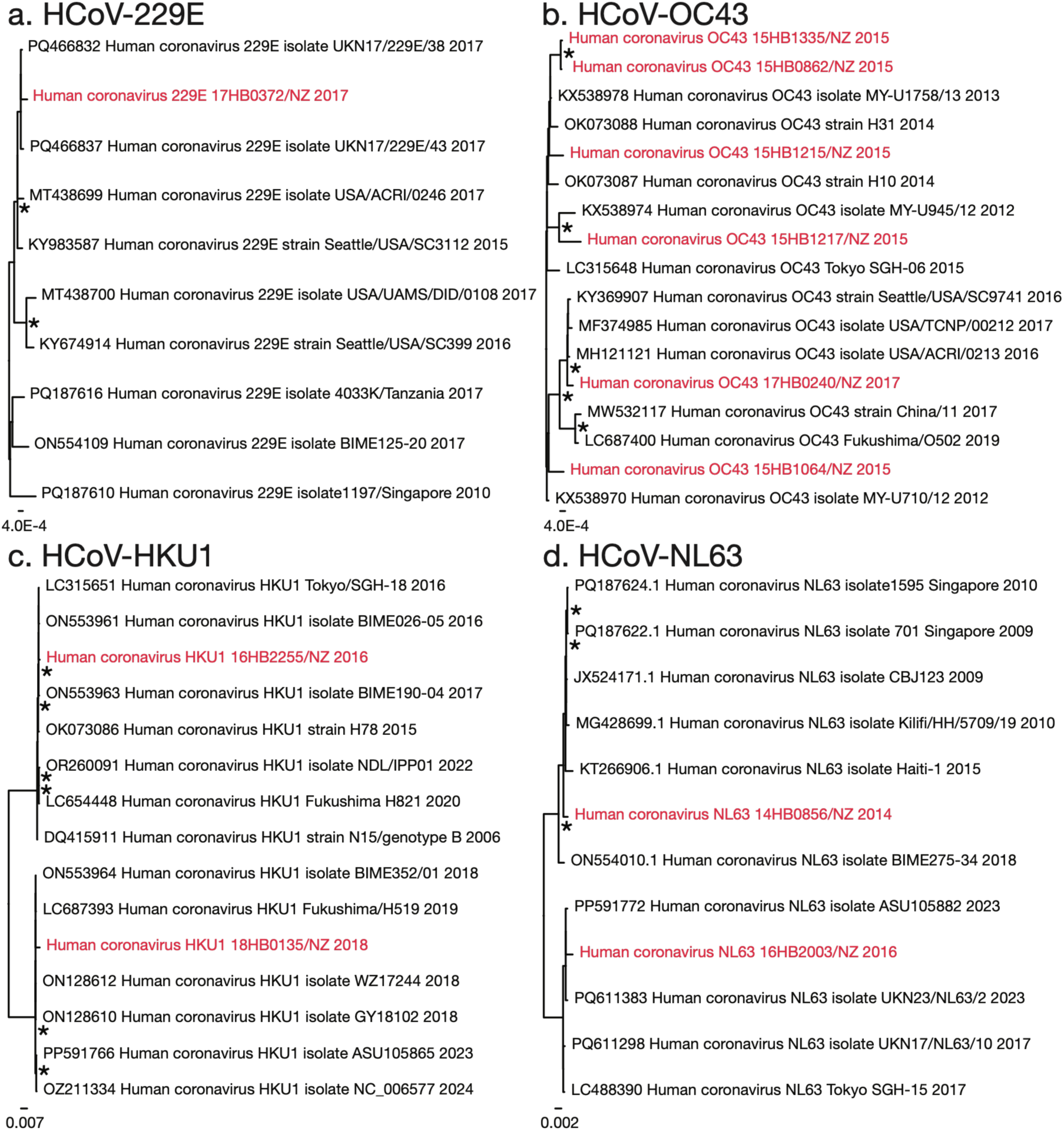
Maximum likelihood phylogenetic trees of Human coronavirus species. Maximum likelihood phylogenetic trees of human coronavirus sequences found in this study (red) and representative nucleotide sequences collected from NCBI taxonomy (black), representing (a) Human coronavirus 229E, (b) Human coronavirus OC43, (c) Human coronavirus HKU1 and (d) Human coronavirus NL63. Phylogenetic trees are midpoint rooted for clarity, and nodes with bootstrap values of >95% are noted with asterisks. Branches are scaled by number of nucleotide substitutions per site shown in the scale bar.

We identified several DNA viruses, including a partial sequence of Human alphaherpesvirus 3 (2,110 nucleotides) (Figure 6). After mapping this sequence (assembled contigs) to a reference genome (NC_001348.1) to improve nucleotide sequence length, we recovered 34,788 nucleotides of the 125kb alphaherpesvirus genome that shared 100% nucleotide similarity with Human alphaherpesvirus 3 isolate Var160 (KC112914) in the envelope glycoprotein (E), previously sampled from a vesicular fluid sample in Mexico in 2007 (46). The envelope glycoprotein (E) is abundantly expressed on the surface of infected cells and is essential for virus replication and cell to cell transmission. We also identified a partial sequence (529 nucleotides) of Epstein-Barr virus (i.e. Human gammaherpesvirus 4) in the same patient sample (17HB0495). Mapping this sequence to a reference genome (NC_007605) we recovered 6,717 nucleotides of the 172kb gammaherpesvirus genome. This segment shared 98% nucleotide similarity with a hypothetical protein of Human gammaherpesvirus 4 isolate HKNPC44 (MH590555.1), previously detected in saliva from nasopharyngeal carcinoma patients in Hong Kong (47). This patient sample was collected in 2017 in Counties Manukau from a female patient of unknown age. We also identified bacterial co-infection in this patient sample, detecting *Streptococcus pneumoniae* at an abundance of 62 RPM.

**Figure 6.**
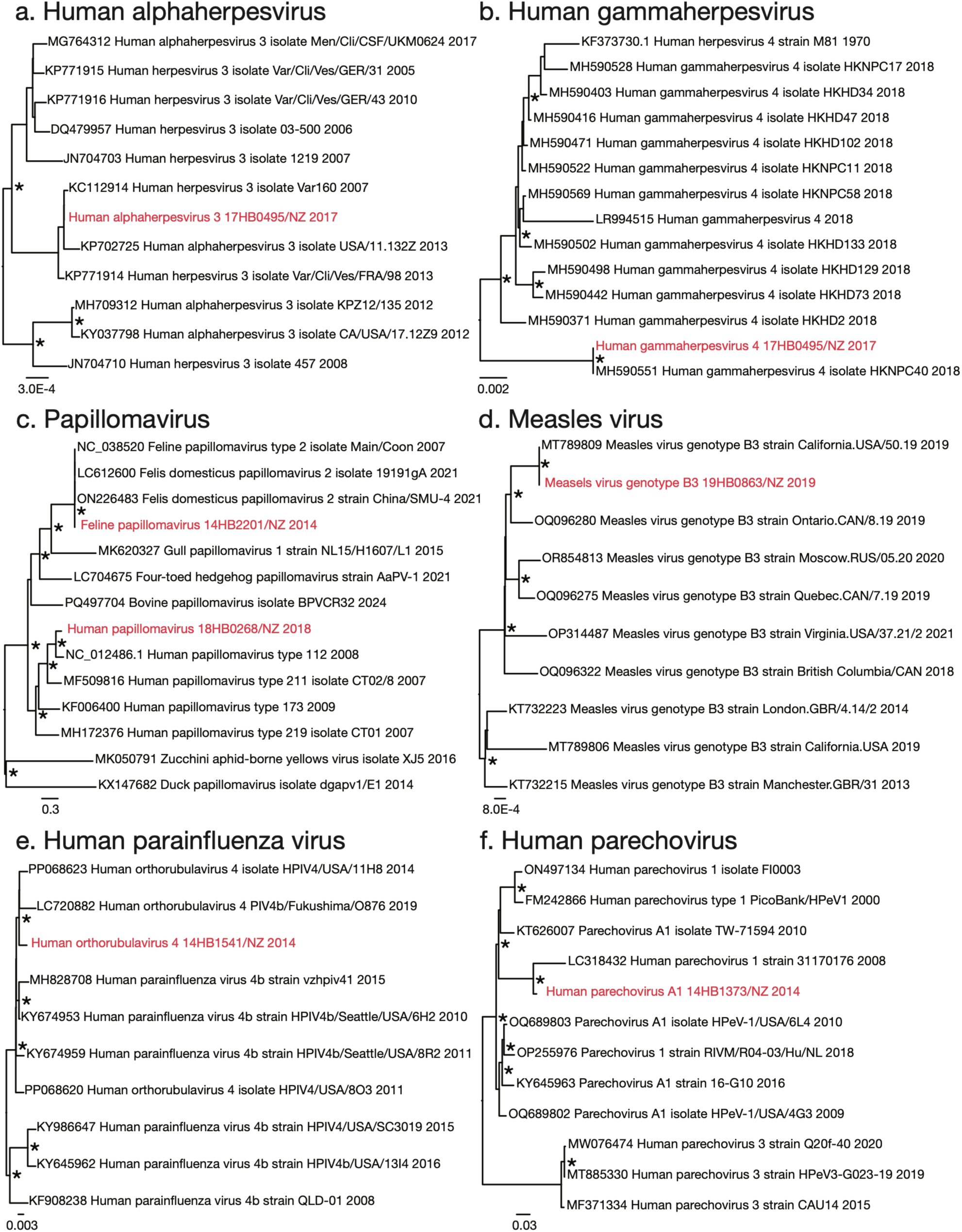
Maximum likelihood phylogenetic trees of identified viral species. Maximum likelihood phylogenetic trees of viral sequences identified here (red) and representative nucleotide sequences collected from NCBI taxonomy (black), representing (a) Human alphaherpesvirus, (b) Human gammaherpesvirus, (c) Papillomavirus, (d) Measles virus, (e) Human parainfluenza virus and (f) Human parechovirus. Phylogenetic trees are midpoint rooted for clarity, and nodes with bootstrap values of >95% are noted with asterisks. Branches are scaled by number of nucleotide substitutions per site shown in the scale bar.

Notably, of the two papillomaviruses identified, one shared high sequence similarity with feline papillomaviruses, including 99% nucleotide similarity with Feline papillomavirus type 2 isolate Main Coon 2007 (NC_038520), identified from a *Felis catus domesticus* (domestic cat) biopsy specimen (48). A partial sequence of the E2 gene (485 bp) was identified. The second papillomavirus shared 83% nucleotide sequence similarity with Human papillomavirus type 112 (NC_012486) in the E2 gene. Upon phylogenetic analysis, these papillomaviruses fell into separate feline and human clades (Figure 6). Given the unexpected host association and limited sequence length of the feline papillomavirus-like sequence, this detection should be interpreted cautiously and could reflect incidental exposure, contamination or sequence misassignment rather than human infection. The two samples in which papillomavirus sequences were detected also exhibited the richest infectomes in the dataset, with more than four microbial species detected in each sample. These included *Staphylococcus aureus*, *Streptococcus pneumoniae*, *Klebsiella pneumoniae*, *Moraxella catarrhalis*, *Haemophilus influenzae* and *Rhinovirus B* (48).

We identified Measles virus genotype B3 NZ 2019, which shared 100% nucleotide similarity with Measles virus genotype B3 strain MVs/California (MT789809) sampled in the same year (Figure 6). Within this sample, we detected co-infection with *Staphylococcus aureus* at relatively low abundance (2 RPM). We identified a full length (17,392 nucleotides) parainfluenza virus genome, Human orthorubulavirus 4 NZ 2014 (Figure 6), which shared 99% similarity to Human orthorubulavirus 4 isolate HPIV4/USA/2014 (PP068623) (49). This sequence was highly abundant (11,863 RPM) and was collected in 2014 from a 64 year old female patient. Finally, we identified a full length (7,298 nucleotides) Human parechovirus A1 NZ 2014 genome (Figure 6), sharing 91% nucleotide similarity with Parechovirus A1 strain 16-G10 (KY645963) identified in USA in 2016. This relatively abundant sequence, detected at 591 RPM, was detected in a nine month old infant, with no other microbial species identified.

### Bacterial and fungal species uncovered

The most commonly identified bacterial species were *Streptococcus pneumoniae* (n = 34), *Pseudomonas aeruginosa* (n = 22), *Staphylococcus aureus* (n = 20) and *Moraxella catarrhalis* (n = 19). Other species were detected at relatively low frequencies, including *Acinetobacter baumannii* (n = 8), *Acinetobacter calcoaceticus* (n = 2), *Klebsiella pneumoniae* (n = 2) and *Haemophilus influenzae* (n = 4). While *Serratia marcescens* was only detected in a single sample (n = 1), it exhibited very high relative abundance (453,377 RPM). Of the 90 individuals in which we detected a bacterial species, 24% were children (0-5 years), 33.3% were aged between 5-65, while 33.3% were aged >65, and 8.9% were of an unknown age. Using all samples (n = 299) Fisher’s exact test indicated that bacterial detection did not differ significantly by age group (p = 0.22) or by sex (p = 0.31). The sampling years contributing the most bacterial detections were 2015 and 2017, accounting for 27% and 30% of detections, respectively. In terms of sample location, 68% of bacterial detections were collected in Auckland and 32% from Counties Manukau.

The most commonly identified fungal species was *Naganishia albida* (n = 9), followed by *Penicillium chrysogenum* (n = 7), *Acremonium sclerotigenum* (n = 2) and *Purpureocillium lilacinum* (n = 2). All four fungal species were detected at high relative abundances ranging from 28,039 to 170,652 RPM. Of the 19 individuals in which we detected a fungal species, 26% were children (0-5 years), 42% were aged 5-65 years, 26% were aged >65, and 5% were of an unknown age. Fungal species were most commonly detected in 2015 (47%), and most cases were detected in Auckland (84%). Females accounted for the majority of cases with a fungal organism detected (68%). Due to very low prevalence and sparse counts in some age categories, statistical comparisons across age groups using Fisher’s test was not possible.

### Factors shaping infectome composition and diversity

We investigated whether the alpha diversity of SARI patient infectomes, which refers to both species richness (the number of distinct species detected) and their relative abundance (evenness), was influenced by host and sampling factors including age, sex, year of collection or geographic location (District Health Board, DHB). We assessed the alpha diversity using richness, Simpson and Shannon indices to capture the overall microbial structure, reflecting species richness and evenness. Abundance was analysed separately to identify specific species that differed in their abundance across patient groups.

When alpha diversity was measured using the Simpson index, we observed significant associations with age (p = 0.02) and year of collection (p = 0.02). The Simpson index places greater weight on the most abundant taxa in a community, making it more sensitive to changes driven by dominant species. In contrast, the Shannon index, which balances both richness and evenness, indicated that infectome composition was influenced only by year of collection (p = 0.001) (Supplementary Figure 2). The difference between 2015 and 2016 (p = 0.001) appeared to drive this pattern, which may in part reflect the greater number of samples collected in 2015.

Using the Kruskal-Wallis rank sum test on microbial abundance, we found that age, year of collection and DHB significantly affected infectome abundance (p = 0.0001, 0.0002 and 0.03 respectively). In contrast, sex was not significantly associated with abundance differences. To identify which specific groups contributed to these differences, we used the Dunn-Bonferroni post-hoc pairwise tests. We found significant differences in alpha diversity across age groups, 0-5 and 5-65 (p = 0.00005) and between 0-5 and 65+ (p = 0.002). Using the Dunn’s test we identified the years driving differences in alpha diversity were comparisons between 2015 and 2016 (p = 0.0003) and 2015 and 2017 (p = 0.008). Pairwise testing found that DHB also significantly influenced alpha diversity (p = 0.03).

To assess whether richness alone contributed to alpha diversity patterns, we applied a generalised linear model and found that age, sex and year of collection were all significant predictors of infectome diversity (Figure 7). In terms of age, richness differed significantly between the 0 – 5 and 5 – 65 year groups (p = 0.001), as well as between the 0 – 5 and >65 groups (p = 0.03). The >65 year group exhibited the highest richness, with one sample containing 5 species. A significant difference in richness was also observed between female and male samples (p = 0.04), with females exhibiting approximately 28% higher richness than males. Year of collection further explained variation in richness, with significant differences detected between 2014 and 2016 (p = 0.02), 2015 and 2016 (p = 0.0007), 2015 and 2017 (p = 0.02), and 2016 and 2018 (p = 0.009) (Supplementary Table 4). Richness peaked in 2018, with one sample containing 5 species. All statistical analyses were performed on the full dataset and on a subset excluding samples with missing metadata (age or sex) and later years (2018–2020) combined due to low sample numbers (see Supplementary Table 4).

**Figure 7.**
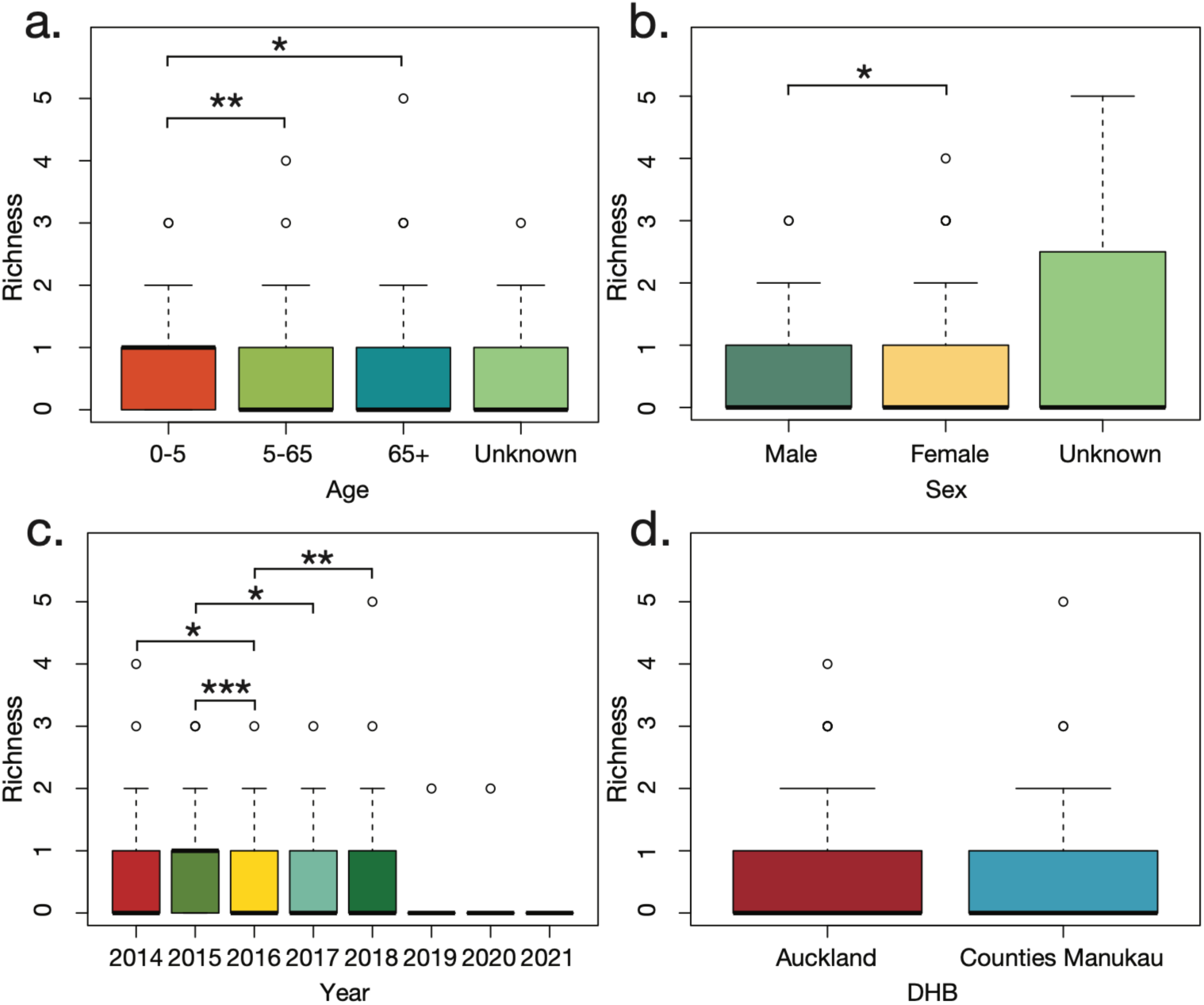
Alpha diversity (Richness). Species richness of infectomes for (a) age (b) sex, (c) year and (d) DHB.

To investigate beta diversity, which measures differences in microbial composition between groups, we grouped together the 10 samples collected between 2018 and 2020 when investigating sampling time. Similarly, we removed 10 samples with missing data (i.e. unknown patient age and one sample with unknown sex). A principal coordinates analysis based on the Bray-Curtis dissimilarity matrix, with ellipses to indicate group clustering, revealed clustering by both year of collection and age group (Figure 8). These observations were supported by PERMANOVA tests, which showed significant clustering by age (p = 0.001) and year (p = 0.02). We found no significant clustering for patient sex (p = 0.2) or DHB (p = 0.3) (Figure 8).

**Figure 8.**
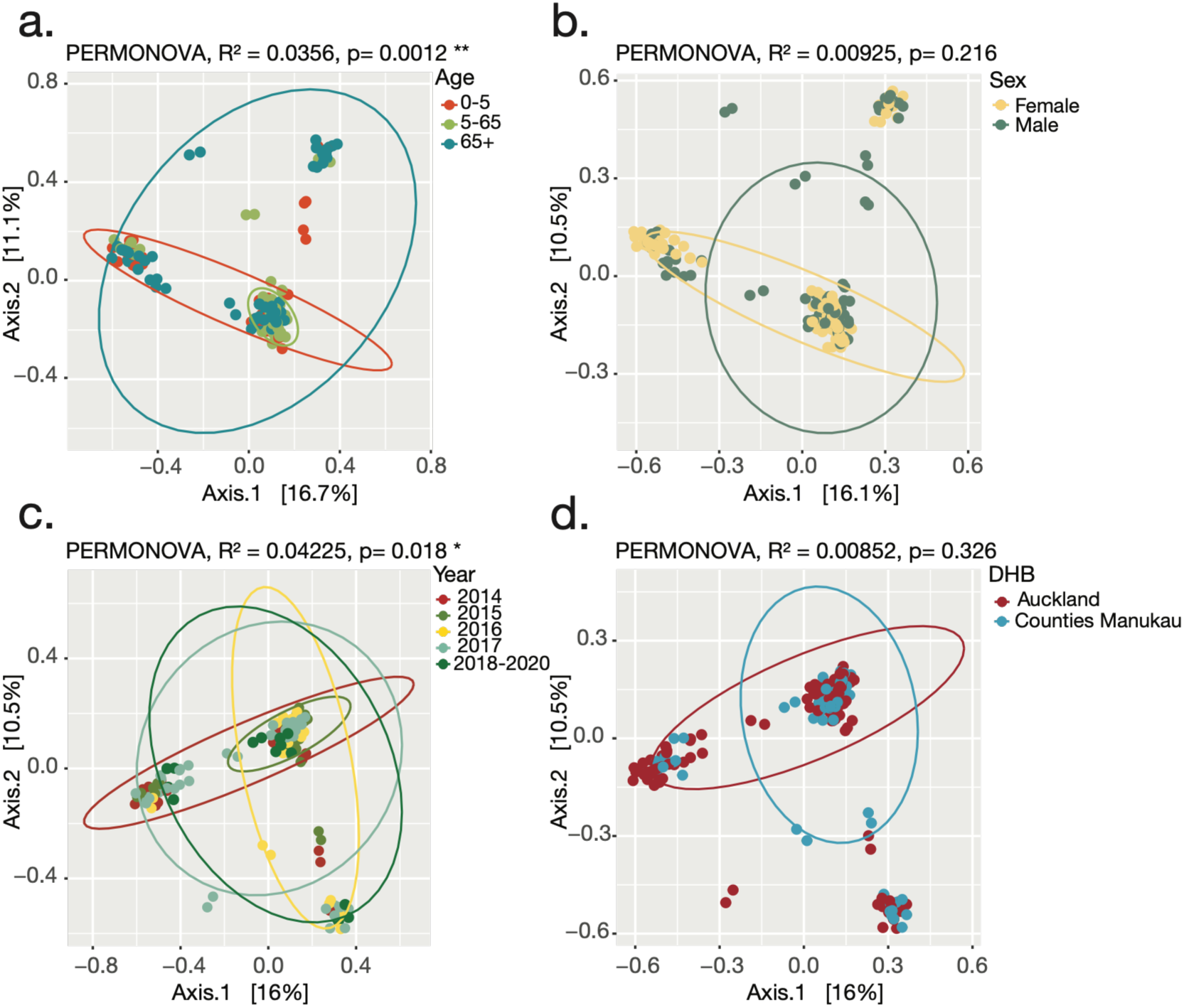
Beta diversity. (a) Principal coordinates analysis showing microbial community clustering according to age (samples with unknown ages removed), (b) sex (samples with unknown sex removed) and (c) years collected (2018-2020 combined due to low sample count) and (d) DHB. Points are jittered for clarity.

### Network analysis of patterns of microbial co-detection

Out of 129 samples, 74% had one microbial species identified, 19% had two species, 5% had three species and 0.8% each had four or five species. There was no significant difference in age among the 33 individuals in which more than one microbial species was detected (Chi-squared test, p = 0.4), with 33% aged 0-5 years, 27% aged 5-65 years, 27% aged >65 years and 12% of unknown age. A chi-squared test was appropriate for assessing co-detection patterns within the 129 positive samples because the contingency tables contained sufficient counts across age groups. Only two patient samples contained more than one viral species, and one sample contained more than one fungal species, whereas 17 samples contained at least two bacterial species. The most common viral-bacterial co-detection was Human rhinovirus C and Moraxella catarrhalis (n = 3), followed by *Streptococcus pneumoniae* and papillomavirus (n = 2). Bacterial-fungal co-detections were detected in three samples, while viral-fungal co-detections occurred in four (Figure 9). The microbial species most frequently found together were *Streptococcus pneumoniae* and *Staphylococcus aureus*, co-occurring in six samples. The second most common pair was Human rhinovirus C and Moraxella catarrhalis, co-occurring in three samples. Most samples with multiple microbial detections were collected in 2015 (39%) and 2017 (21%). Of the 33 samples with multiple microbial detections, there was no difference among female (54%) and male (42.4%) patients (p = 0.5). Most samples that had multiple microbial detections were collected in Auckland (61%) with the remainder from Counties Manukau (39%), however, the difference in co-detection rates between DHBs was not significant (Chi-squared test with Yates’ correction, p = 0.14).

**Figure 9.**
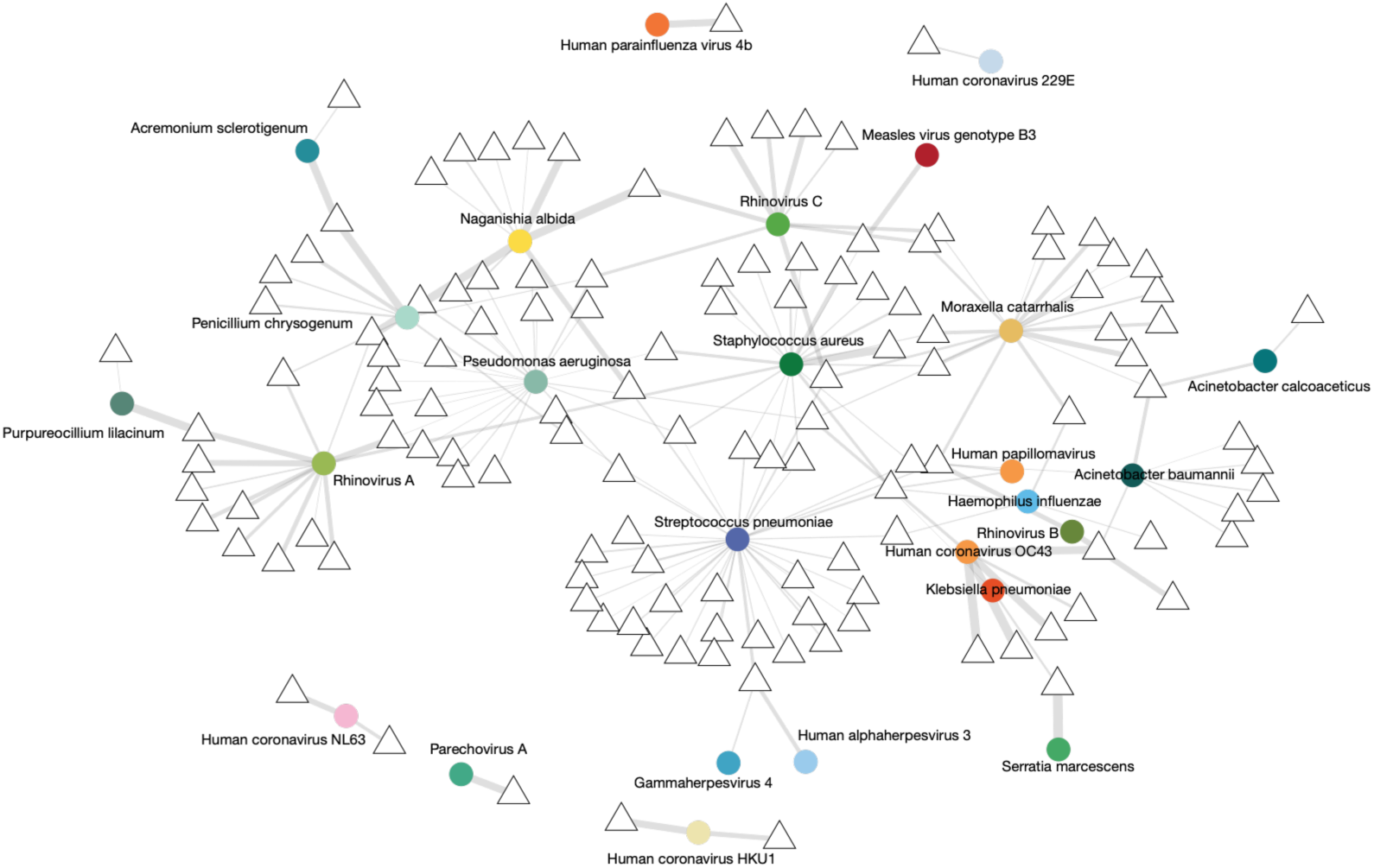
Bipartite network Illustrating the connections between microbial species identified in this study (coloured circles) and SARI patients (white triangles). Branch thickness is weighted by the standardised abundance (in RPM) of microbial transcripts. Patients in which no microbial species were identified are not shown.

To describe patterns of microbial co-occurrence, a bipartite network was constructed based on microbe-patient associations (Figure 9). The resulting network included 155 nodes (26 microbial species and 129 patient samples) and 174 edges, where each edge represented detection of a microbial species in a patient sample. The network exhibited low connectance (5.2%) and low-moderate Nestedness as a measure of Overlap and Decreasing Fill (NODF = 4.3). Interaction Shannon diversity was 2.21 and interaction evenness was 0.27, reflecting the predominance of a small number of frequently detected species. Linkage density was low (1.5), and most samples were associated with only one or two species (mean degree = 1.4). Species nodes had a mean degree of 6.7 and a maximum of 34, reflecting the frequent detection of *Streptococcus pneumoniae* (n = 34). Community detection using the Louvain algorithm identified 14 modules (modularity = 0.71). In the absence of formal null-model testing, these network properties and modules are presented descriptively and should not be interpreted as evidence of specific ecological interactions between microbial species.

### Determining virulence factors in bacterial species

We examined transcripts corresponding to genes with previously characterised virulence-associated functions in the four most abundant bacterial species identified across all patient samples. These sequence-based assignments indicate expression of genes with reported virulence-associated functions but do not demonstrate phenotypic virulence or establish that the detected organism contributed to disease in the sampled patient. For *Moraxella catarrhalis*, we identified 796 genes with an abundance greater than 1 RPM, out of 1,839 genes in the reference genome (Figure 10). The most abundantly expressed gene was an OmpA family protein, a well-characterised virulence factor in Gram-negative bacteria (50,51). OmpA proteins are involved in host cell adhesion, evasion of immune responses and intracellular survival (52). These proteins are commonly associated with respiratory tract infections (52). We also detected a relatively highly expressed TonB-dependent receptor protein, which has previously been linked to macrolide resistance in M. catarrhalis (53). Additionally, we identified an alkyl hydroperoxide reductase system C, which supports bacterial survival under oxidative stress conditions (50–54).

**Figure 10.**
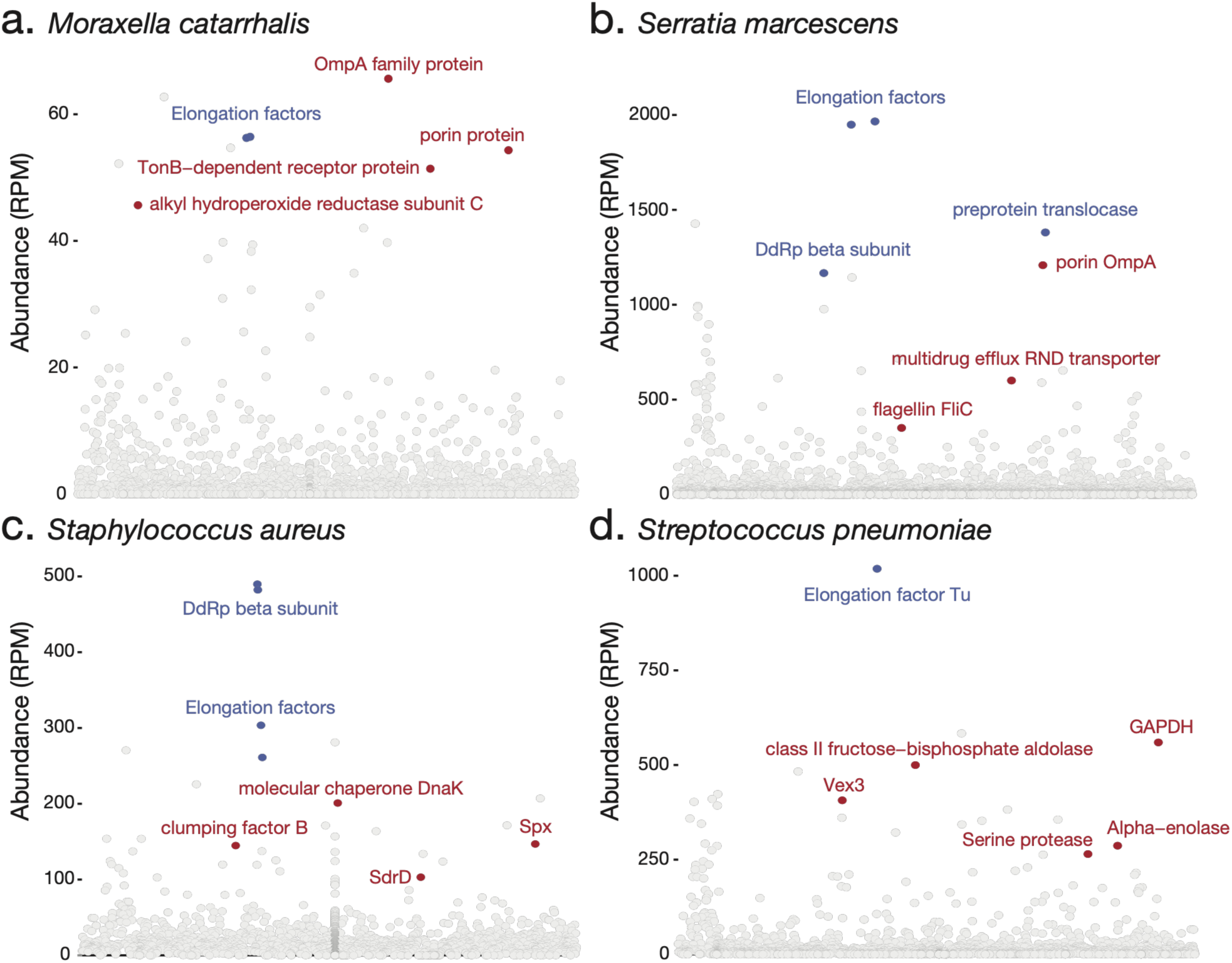
Virulence gene expression. Gene expression of most abundant bacterial species in reads per million (RPM). Red labels highlight highly expressed known virulence genes, while blue labels indicate highly expressed essential genes.

For *Serratia marcescens*, 3,184 genes were identified with an abundance greater than 1 RPM out of a total of 4,781 genes in the reference genome. Among these we detected a highly expressed OmpA outer membrane protein with porin activity, which has been shown to contribute to structural integrity and ion permeability, aiding survival under osmotic stress conditions in *S. marcescens*. We also identified a highly expressed resistance-nodulation division superfamily efflux system, a key contributor to multidrug resistance in *S. marcescens* (55–56).

In *Staphylococcus aureus*, we identified 2,369 genes with an abundance greater than 1 RPM, out of 2,767 genes in the reference genome. Among these, we detected several known virulence factors that were highly expressed. The serine-aspartate repeat-containing protein D (SdrD) was identified at relatively high abundance; this adhesin facilitates binding to human nasal squamous epithelial cells and has been implicated in nasal colonization by *S. aureus*. We also detected Spx, a global stress response regulator involved in biofilm formation and cellular growth. Spx has been previously shown to contribute to *S. aureus* persistence by regulating oxidative stress responses and influencing biofilm formation dynamics. Another highly abundant gene was DnaK, a molecular chaperone known to facilitate adhesion to eukaryotic cells and support biofilm development. Loss of DnaK has been associated with significantly reduced virulence in S. aureus. Lastly, we identified high expression of clumping factor B (ClfB), a surface protein that promotes adhesion to host epithelial cells and has been directly linked to efficient nasal colonization (57–60).

In *Streptococcus pneumoniae*, 1,365 genes were identified with an abundance greater than 1 RPM, out of 2,116 genes in the reference genome. Among the highly expressed genes was a class II fructose-bisphosphate aldolase, which has been previously characterised as a bacterial adhesin that contributes to host colonization by promoting attachment to epithelial cells. We also identified glyceraldehyde-3-phosphate dehydrogenase (GAPDH) and alpha enolase, both of which are known to bind plasminogen and facilitate bacterial translocation across mucosal barriers, supporting dissemination during infection. We also detected serine protease PrtA, a virulence-associated protein involved in the cleavage of host proteins such as immunoglobulins, complement components and extracellular matrix proteins. Deletion of PrtA has been shown to significantly reduce pneumococcal virulence in murine models. Additionally, an ATP-binding cassette (ABC) transporter permease subunit Vex3 was identified. ABC transporters in S. pneumoniae have been associated with antibiotic resistance, including vancomycin tolerance. Specifically, the vex123 operon has been linked to increased resistance with vex3 mutants demonstrating greater antibiotic tolerance than wild type strains (61–65).

### Uncovering the resistome

Sequence-based detection of antimicrobial resistance genes indicates the presence and transcription of sequences homologous to characterised resistance determinants, but does not demonstrate phenotypic antimicrobial resistance without isolate-based susceptibility testing. We identified 438 ARGs with an abundance greater than 1 RPM (Figure 11). From these, the top 100 most abundant (i.e. most expressed) ARGs were selected, ranging from 66 to 17,724 RPM (average 538 RPM). Six highly abundant ARGs (blaOXA-60d_1_AY664506, blaOXA-22_1_AF064820, blaOXA-444_1_CP010800, blaOXA-443_1_LC030178, blaOXA-60c_1_AY664505, blaOXA-60b_1_AY664504) were removed due to suspected contamination, as they were detected in every sample. The number of ARGs detected per sample ranged from 0 to 19, with an average of 4.4 ARGs per sample (66).

**Figure 11.**
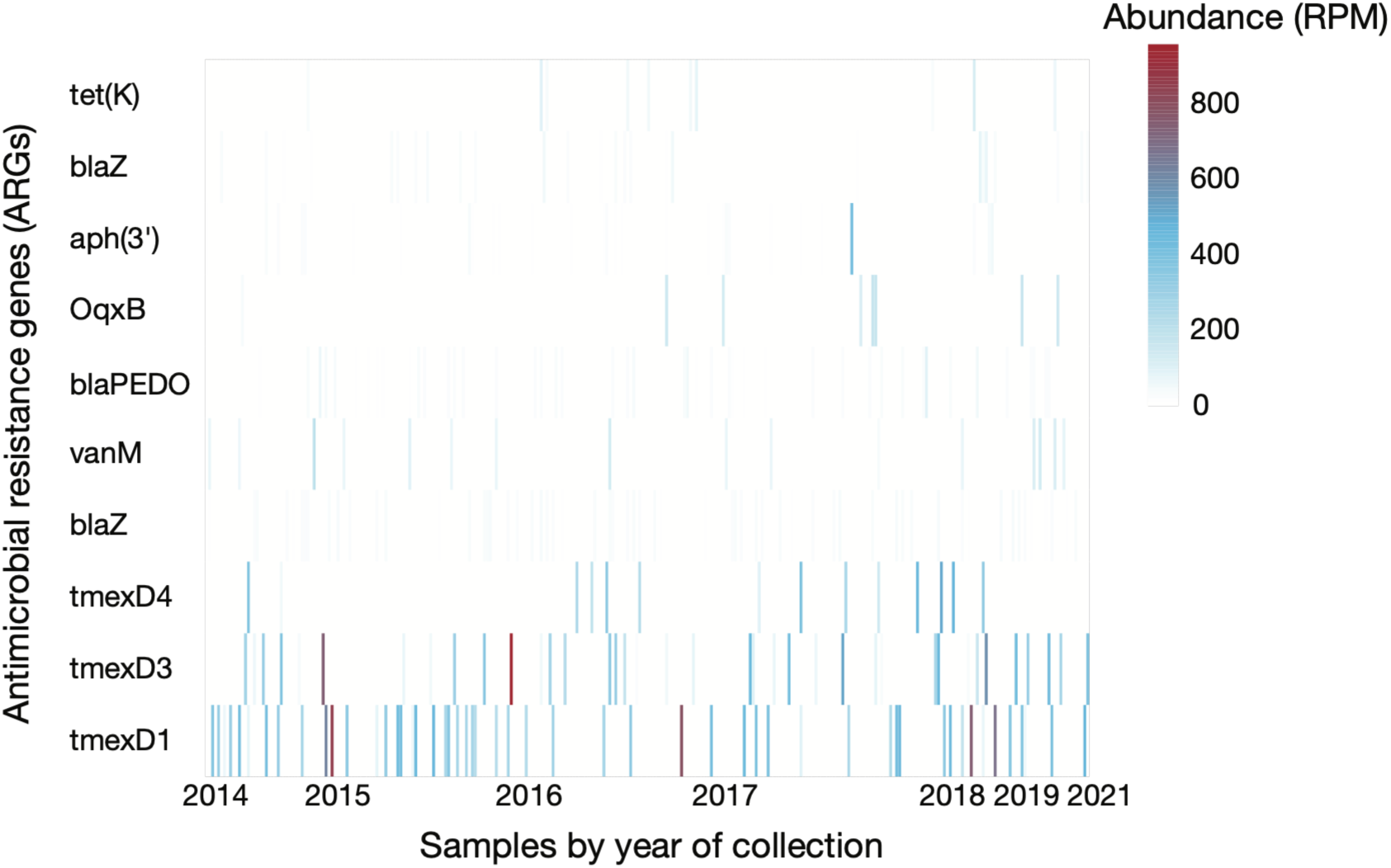
Heatmap of antimicrobial resistance genes. The 10 most abundant antimicrobial resistance genes (ARGs) identified across all (299) patient samples, across sampling time. The colour corresponds to the abundance of reads, expressed as reads per million (RPM). Samples are ordered by time along the x-axis.

The tmexD1_1_MK347425 sequence was identified in Klebsiella pneumoniae in chickens (Figure 11). It was the most abundant sequence in the dataset (with 17,724 RPM) and the second most frequently detected sequence (52 samples). This sequence encodes a resistance–nodulation– division efflux pump gene cluster located on a plasmid, which confers resistance to multiple antibiotic classes, including tetracyclines, quinolones, cephalosporins, and aminoglycosides (67). The blaZ_3_CP000732 sequence was identified in paediatric cases of Staphylococcus aureus including both methicillin resistant and methicillin susceptible strains. It was the most frequently detected sequence (69 samples) and was found in high abundance (1,697 RPM). This plasmid-encoded gene encodes a class A β-lactamase, an important mechanism of β-lactam resistance (68). mecA, the methicillin-resistance gene, was not detected in any sample. The tmexD3_1_LC633285 sequence was identified in Klebsiella aerogenes and encodes a plasmid associated RND multidrug efflux transporter permease subunit. It was the second most abundant sequence identified in this dataset (10,135 RPM), detected in 39 samples. The tmexD4_1_CP091084 sequence was identified in Enterobacter roggenkampii and encodes a plasmid associated RND multidrug efflux transporter permease subunit (69). It was found in 14 samples and at high abundance (4,044 RPM). The OqxB_1_EU370913 sequence, a member of the RND efflux pump family, was identified in 8 patient samples.

## Discussion

This study provides an overview of microbial communities present in nasopharyngeal samples from patients with severe acute respiratory infection (SARI) in Aotearoa New Zealand. Using an unbiased metatranscriptomic approach with a focus on taxa with recognised or potential associations with respiratory infection, we characterised transcriptionally active viral, bacterial and fungal communities in 299 patients who tested negative for common respiratory viruses by PCR. The approach also captured transcripts corresponding to bacterial virulence-associated factors and antimicrobial resistance genes, providing functional information in addition to taxonomic identification.

This approach enabled the detection of a wide spectrum of viruses, including species commonly associated with respiratory infections as well as less frequently screened microbes (70). Several viral species not included in the routine testing applied to these samples, including seasonal human coronaviruses, parainfluenza virus 4 and human parechovirus A1, were identified. Rhinoviruses were detected in 23 patients despite not being identified during the original PCR screening. Alignment of the assay primers against the available rhinovirus sequences revealed perfect or near-perfect complementarity, suggesting that primer mismatch alone is unlikely to explain the discordant results (71). However, the reason for the discrepancy between the original PCR results and subsequent metatranscriptomic detection cannot be determined retrospectively. Potential explanations include differences in template abundance, RNA integrity, assay sensitivity, amplification efficiency or other technical factors. These detections should therefore be interpreted as discordance between the two approaches rather than confirmed false-negative PCR results. Detection of both viruses included in the original diagnostic panel and viruses not routinely screened for demonstrates the complementary information that metatranscriptomic sequencing can provide when investigating diagnostically unresolved respiratory infections (70,71).

We also identified bacterial and fungal species with recognised or potential associations with respiratory infection. *Streptococcus pneumoniae*, *Staphylococcus aureus* and *Moraxella catarrhalis* were among the most frequently detected bacterial species, and transcripts corresponding to characterised virulence-associated and antimicrobial resistance genes were identified (72,73). These sequence-based findings do not establish phenotypic virulence, antimicrobial resistance or a causal role in SARI. The fungal taxa included species typically considered environmental or opportunistic, and their detection in nasopharyngeal samples may represent colonisation, incidental exposure or infection. *Naganishia albida* was detected in nine patients, including three high-abundance detections. *N. albida* is a rare opportunistic pathogen that has been reported in patients with respiratory symptoms such as cough and fever (74,75). In a clinical case in China, NGS-based detection contributed to identification of N. albida and subsequent clinical management (75). This example illustrates the potential value of sequencing for identifying uncommon organisms, while the clinical significance of the detections in our cohort cannot be established from the available data (72–75).

Multiple microbial species were detected in a subset of samples, including Rhinovirus C with *Moraxella catarrhalis* and Rhinovirus A with *S. aureus*. Interactions between these organisms have previously been associated with altered bacterial or viral loads, host responses and disease severity (76–78), although co-detection in the present study does not demonstrate such interactions. Other co-occurrences included Human coronavirus OC43 with opportunistic bacteria such as *Acinetobacter baumannii* and *Pseudomonas aeruginosa*. Similar virus-bacterium interactions have been reported in other respiratory infections, including SARS-CoV-2 and *A. baumannii* (79–83), but whether these organisms interacted or jointly influenced disease in the present cohort cannot be determined.

Nasopharyngeal samples characterise microbial communities of the upper respiratory tract and cannot establish the microbial composition of the lower respiratory tract. Consequently, detection of an organism in these samples should not be interpreted as evidence that it was responsible for a patient’s lower respiratory disease. The absence of a control group (e.g. healthy individuals) further restricted our capacity to distinguish disease-associated microbes from background colonisers (18,70). Without a defined baseline ‘normal’ respiratory microbiota, we relied on published studies to establish commonly detected bacterial and fungal taxa (18,70,84–88). As RNA sequence detection does not necessarily indicate causation (89–92), particularly in polymicrobial contexts, the extent to which the organisms identified directly contributed to SARI symptoms remains uncertain. The retrospective sample set was also temporally uneven, with a large proportion of samples collected in 2015 and relatively few available from later years, particularly during 2020-2021. This sampling imbalance limits interpretation of temporal differences in infectome composition and means that the cohort should not be considered representative of the national SARI burden over the study period. In addition, changes in diagnostic testing during the study period may have introduced ascertainment bias. The introduction of SARS-CoV-2 testing during the COVID-19 pandemic expanded respiratory diagnostic panels and could potentially influence patterns of pathogen detection over time. Nevertheless, the observed reduction in alpha diversity preceded the emergence of SARS-CoV-2 and associated lockdowns, suggesting this factor is unlikely to fully explain the temporal patterns observed.

The current SARI surveillance system in New Zealand provides valuable insights but remains geographically limited and resource intensive, restricting its scalability across the wider healthcare system (13). Our findings illustrate how targeted PCR and metatranscriptomic sequencing provide different but complementary information. PCR offers rapid and sensitive detection of predefined targets, whereas unbiased sequencing can detect organisms outside those targets and provide genomic information for surveillance (71,93). Practical implementation of metatranscriptomics in routine diagnostics would require consideration of cost, turnaround time, bioinformatics expertise, contamination control and the clinical interpretation of polymicrobial findings. Further evaluation of genomics-informed approaches could strengthen investigation of diagnostically unresolved cases and support surveillance and response to emerging infectious threats (94–96).

This study presents the first metatranscriptomic characterisation of PCR-negative SARI samples in Aotearoa New Zealand. By sequencing 300 samples collected between 2014 and 2021, we identified 13 viral, nine bacterial and four fungal species, including well-established respiratory pathogens as well as less frequently detected or unexpected taxa. The detection of multiple microbial species, antimicrobial resistance gene transcripts and virulence-associated gene transcripts highlights the breadth of information obtainable from metatranscriptomic sequencing. However, these findings should be interpreted as microbial and functional signatures present in nasopharyngeal samples rather than direct evidence of disease causation or phenotypic antimicrobial resistance.

In the context of routine diagnostics in New Zealand, testing strategies are guided by factors including cost, throughput and turnaround time, with high-throughput RT-PCR assays commonly employed. Our findings identify additional organisms and microbial combinations present in PCR-negative SARI samples, including viruses outside the diagnostic targets applied during the study period. Such information could help inform future evaluation and refinement of respiratory testing panels and surveillance strategies. Metatranscriptomics is unlikely to replace targeted PCR for routine high-throughput testing, but may provide a useful complementary approach for selected diagnostically unresolved cases, surveillance and investigation of unexpected or emerging respiratory pathogens.

## Data availability

Genomic data generated in this study are available on NCBI’s GenBank and SRA database (BioProject: PRJNA1437773).

## Acknowledgements

SHIVERS-I Science Management Group: Michael G. Baker, Nikki Turner, Cameron C. Grant, Colin McArthur, Sally Roberts, Adrian Trenholme, Conroy Wong, Paul Thomas, Richard Webby, Mark G. Thompson, and Marc-Alain Widdowson; The research nurses at Auckland District Health Board (ADHB): Alicia Stanley, Kathryn Haven, Bhamita Chand, Pamela Muponisi, Debbie Aley, Claire Sherring, Miriam Rea, Judith Barry, Tracey Bushell, Julianne Brewer, Catherine McClymont; The research nurses at Counties Manukau District Health Board (CMDHB): Shirley Lawrence, Shona Chamberlin, Reniza Ongcoy, Kirstin Davey, Emilina Jasmat, Maree Dickson, Annette Western, Olive Lai, Sheila Fowlie, Faasoa Aupa’au, Louise Robertson; The New Zealand Institute of Public Health and Forensic Science (PHF Science): Tim Wood, Nayyereh Aminisani, Ruth Seeds, Jacqui Ralston, Judy Bocacao, Wendy Gunn, Andrea McNeill, Sarah Jefferies, Andy Anglemyer; The ADHB Laboratory: Gary McAuliffe, Fahimeh Rahnama; The CMDHB Laboratory: Susan Taylor, Helen Qiao, Fifi Tse, Mahtab Zibaei, Tirzah Korrapadu, Louise Optland, Cecilia Dela Cruz;

## Supplementary Information

Table S1. Summary of detected microbial species, number of libraries, and abundance.

Table S2. Summary of samples with multiple microbial detections.

Table S3. Data surrounding the viral sequences identified in this study and the top BLASTn genetic matches.

Table S4. Summary of infectome statistics.

Figure S1. Rhinovirus sequence with annotated PCR primer binding regions illustrating the expected amplicon location.

Figure S2. Box plots showing the mean and interquartile ranges of Shannon diversity of infectomes for (a) age, (b) sex, (c) years, and (d) DHB.

## References

1. Das S, Dunbar S, Tang YW. Laboratory diagnosis of respiratory tract infections in children – the state of the art. Front Microbiol. 2018;9:2478.

2. Barenfanger J, Drake C, Leon N, Mueller T, Troutt T. Clinical and financial benefits of rapid detection of respiratory viruses: an outcomes study. J Clin Microbiol. 2000;38(8):2824–8.

3. Byington CL, Castillo H, Gerber K, Daly JA, Brimley LA, Adams S, et al. The effect of rapid respiratory viral diagnostic testing on antibiotic use in a children’s hospital. Arch Pediatr Adolesc Med. 2002;156(12):1230.

4. Akers IE, Weber R, Sax H, Böni J, Trkola A, Kuster SP. Influence of time to diagnosis of severe influenza on antibiotic use, length of stay, isolation precautions, and mortality: a retrospective study. Influenza Other Respir Viruses. 2017;11(4):337–44.

5. El Kholy AA, Mostafa NA, Ali AA, Soliman MMS, El-Sherbini SA, Ismail RI, et al. The use of multiplex PCR for the diagnosis of viral severe acute respiratory infection in children: a high rate of co-detection during the winter season. Eur J Clin Microbiol Infect Dis. 2016;35(10):1607–13.

6. Babady NE. The FilmArray respiratory panel: an automated, broadly multiplexed molecular test for the rapid and accurate detection of respiratory pathogens. Expert Rev Mol Diagn. 2013;13(8):779–88.

7. Artika IM, Dewi YP, Nainggolan IM, Siregar JE, Antonjaya U. Real-time polymerase chain reaction: current techniques, applications, and role in COVID-19 diagnosis. Genes (Basel). 2022;13(12):2387.

8. Liu Q, Jin X, Cheng J, Zhou H, Zhang Y, Dai Y. Advances in the application of molecular diagnostic techniques for the detection of infectious disease pathogens (review). Mol Med Rep. 2023;27(5):104.

9. Orle KA, Gates CA, Martin DH, Body BA, Weiss JB. Simultaneous PCR detection of Haemophilus ducreyi, Treponema pallidum, and herpes simplex virus types 1 and 2 from genital ulcers. J Clin Microbiol. 1996;34(1):49–54.

10. Kimura M, Miyake H, Kim HS, Tanabe M, Arai M, Kawai S, et al. Species-specific PCR detection of malaria parasites by microtiter plate hybridization: clinical study with malaria patients. J Clin Microbiol. 1995;33(9):2342–6.

11. Alsharksi AN, Sirekbasan S, Gürkök-Tan T, Mustapha A. From tradition to innovation: diverse molecular techniques in the fight against infectious diseases. Diagnostics. 2024;14(24):2876.

12. Garibyan L, Avashia N. Polymerase chain reaction. J Invest Dermatol. 2013;133(3):1–4.

13. Ministry of Health. Acute respiratory illness surveillance report 2023. Client report no. FW24007. Wellington (NZ): Ministry of Health; 2023.

14. Cheung IMY, Paynter J, Broderick D, Trenholme A, Byrnes CA, Grant CC, et al. Severe acute respiratory infection (SARI) due to influenza in post-COVID resurgence: disproportionate impact on older Māori and Pacific peoples. Influenza Other Respir Viruses. 2024;18(11).

15. Huang QS, Baker M, McArthur C, Roberts S, Williamson D, Grant C, et al. Implementing hospital-based surveillance for severe acute respiratory infections caused by influenza and other respiratory pathogens in New Zealand. West Pac Surveill Response J. 2014;5(2):23–30.

16. Yang J, Yang F, Ren L, Xiong Z, Wu Z, Dong J, et al. Unbiased parallel detection of viral pathogens in clinical samples by use of a metagenomic approach. J Clin Microbiol. 2011;49(10):3463–9.

17. Satam H, Joshi K, Mangrolia U, Waghoo S, Zaidi G, Rawool S, et al. Next-generation sequencing technology: current trends and advancements. Biology (Basel). 2023;12(7):997.

18. Li CX, Li W, Zhou J, Zhang B, Feng Y, Xu CP, et al. High-resolution metagenomic characterization of complex infectomes in paediatric acute respiratory infection. Sci Rep. 2020;10(1):3963.

19. World Health Organization. Global epidemiological surveillance standards for influenza. Geneva: World Health Organization; 2013 [cited 2025 Nov 24]. Available from: https://www.who.int/publications/i/item/9789241506601

20. Shu B, Wu KH, Emery S, Villanueva J, Johnson R, Guthrie E, et al. Design and performance of the CDC real-time reverse transcriptase PCR swine flu panel for detection of 2009 A(H1N1) pandemic influenza virus. J Clin Microbiol. 2011;49(7):2614–9.

21. Kim C, Ahmed JA, Eidex RB, Nyoka R, Waiboci LW, Erdman D, et al. Comparison of nasopharyngeal and oropharyngeal swabs for the diagnosis of eight respiratory viruses by real-time reverse transcription-PCR assays. PLoS One. 2011;6(6).

22. Bolger AM, Lohse M, Usadel B. Trimmomatic: a flexible trimmer for Illumina sequence data. Bioinformatics. 2014;30(15):2114–20.

23. Andrews S. FastQC: a quality control tool for high throughput sequence data. Cambridge (UK): Babraham Bioinformatics; 2010 [cited 2025 Mar 10]. Available from: http://www.bioinformatics.babraham.ac.uk/projects/fastqc/

24. Langmead B, Salzberg SL. Fast gapped-read alignment with Bowtie 2. Nat Methods. 2012;9(4):357–9.

25. Li D, Liu CM, Luo R, Sadakane K, Lam TW. MEGAHIT: an ultra-fast single-node solution for large and complex metagenomics assembly via succinct de Bruijn graph. Bioinformatics. 2015;31(10):1674–6.

26. Buchfink B, Reuter K, Drost HG. Sensitive protein alignments at tree-of-life scale using DIAMOND. Nat Methods. 2021;18:366–8.

27. Chang WS, Harvey E, Mahar JE, Firth C, Shi M, Simon-Loriere E, et al. Improving the reporting of metagenomic virome-scale data. Commun Biol. 2024;7(1):1687.

28. Katoh K. MAFFT: a novel method for rapid multiple sequence alignment based on fast Fourier transform. Nucleic Acids Res. 2002;30(14):3059–66.

29. Kroneman A, Vennema H, Deforche K, van der Avoort H, Peñaranda S, Oberste MS, et al. An automated genotyping tool for enteroviruses and noroviruses. J Clin Virol. 2011;51(2):121–5.

30. Nguyen LT, Schmidt HA, von Haeseler A, Minh BQ. IQ-TREE: a fast and effective stochastic algorithm for estimating maximum-likelihood phylogenies. Mol Biol Evol. 2015;32(1):268–74.

31. Kalyaanamoorthy S, Minh BQ, Wong TKF, von Haeseler A, Jermiin LS. ModelFinder: fast model selection for accurate phylogenetic estimates. Nat Methods. 2017;14(6):587–9.

32. Rambaut A. FigTree. Edinburgh: Institute of Evolutionary Biology, University of Edinburgh; 2018 [cited 2026 Oct]. Available from: http://tree.bio.ed.ac.uk/software/figtree/

33. SnapGene. SnapGene software. Chicago (IL): Dotmatics; 2021 [cited 2025 Mar 10]. Available from: https://support.snapgene.com/hc/en-us

34. Marcelino VR, Clausen PTLC, Buchmann JP, Wille M, Iredell JR, Meyer W, et al. CCMetagen: comprehensive and accurate identification of eukaryotes and prokaryotes in metagenomic data. Genome Biol. 2020;21(1):103.

35. Clausen PTLC, Aarestrup FM, Lund O. Rapid and precise alignment of raw reads against redundant databases with KMA. BMC Bioinformatics. 2018;19(1):307.

36. Sprent P. Fisher exact test. In: International encyclopedia of statistical science. Berlin: Springer; 2011. p. 524–5.

37. McHugh ML. The chi-square test of independence. Biochem Med (Zagreb). 2013;23(2):143–9.

38. Kruskal WH, Wallis WA. Use of ranks in one-criterion variance analysis. J Am Stat Assoc. 1952;47(260):583–621.

39. Dunn OJ. Multiple comparisons using rank sums. Technometrics. 1964;6(3):241–52.

40. Ordak M. Multiple comparisons and effect size: statistical recommendations for authors planning to submit an article to Allergy. Allergy. 2023;78(5):1145–7.

41. McMurdie PJ, Holmes S. phyloseq: an R package for reproducible interactive analysis and graphics of microbiome census data. PLoS One. 2013;8(4).

42. Oksanen J, Blanchet FG, Friendly M, Kindt R, Legendre P, McGlinn D, et al. vegan: community ecology package. R package version 2.5–7. 2020.

43. Liao Y, Smyth GK, Shi W. The R package Rsubread is easier, faster, cheaper and better for alignment and quantification of RNA sequencing reads. Nucleic Acids Res. 2019;47(8).

44. Chen L. VFDB: a reference database for bacterial virulence factors. Nucleic Acids Res. 2005;33(Database issue):D325–8.

45. Bortolaia V, Kaas RS, Ruppe E, Roberts MC, Schwarz S, Cattoir V, et al. ResFinder 4.0 for predictions of phenotypes from genotypes. J Antimicrob Chemother. 2020;75(12):3491–500.

46. Garcés-Ayala F, Rodríguez-Castillo A, Ortiz-Alcántara JM, Gonzalez-Durán E, Segura-Candelas JM, Pérez-Agüeros SI, et al. Full-genome sequence of a novel varicella-zoster virus clade isolated in Mexico. Genome Announc. 2015;3(4).

47. Hui KF, Chan TF, Yang W, Shen JJ, Lam KP, Kwok H, et al. High-risk Epstein–Barr virus variants characterized by distinct polymorphisms in the EBER locus are strongly associated with nasopharyngeal carcinoma. Int J Cancer. 2019;144(12):3031–42.

48. Lange CE, Tobler K, Markau T, Alhaidari Z, Bornand V, Stöckli R, et al. Sequence and classification of FdPV2, a papillomavirus isolated from feline Bowenoid in situ carcinomas. Vet Microbiol. 2009;137(1–2):60–5.

49. Goya S, Greninger AL. Distinct evolutionary signatures of human parainfluenza viruses 2 and 4 reveal host antagonism divergence and phylogenetic discordance. Mol Biol Evol. 2025;42(10).

50. Smith SGJ, Mahon V, Lambert MA, Fagan RP. A molecular Swiss army knife: OmpA structure, function and expression. FEMS Microbiol Lett. 2007;273(1):1–11.

51. Vila-Farrés X, Parra-Millán R, Sánchez-Encinales V, Varese M, Ayerbe-Algaba R, Bayó N, et al. Combating virulence of Gram-negative bacilli by OmpA inhibition. Sci Rep. 2017;7(1):14683.

52. Confer AW, Ayalew S. The OmpA family of proteins: roles in bacterial pathogenesis and immunity. Vet Microbiol. 2013;163(3–4):207–22.

53. Zhang Z, Yang Z, Xiang X, Liao P, Niu C. Mutation of TonB-dependent receptor encoding gene MCR_0492 potentially associates with macrolide resistance in Moraxella catarrhalis isolates. Infect Drug Resist. 2022;15:2419–26.

54. Hoopman TC, Liu W, Joslin SN, Pybus C, Brautigam CA, Hansen EJ. Identification of gene products involved in the oxidative stress response of Moraxella catarrhalis. Infect Immun. 2011;79(2):745–55.

55. Gangadharappa BS, Rajashekarappa S, Sathe G. Proteomic profiling of Serratia marcescens by high-resolution mass spectrometry. BioImpacts. 2020;10(2):123–35.

56. Toba S, Minato Y, Kondo Y, Hoshikawa K, Minagawa S, Komaki S, et al. Comprehensive analysis of resistance-nodulation-cell division superfamily (RND) efflux pumps from Serratia marcescens Db10. Sci Rep. 2019;9(1):4854.

57. Askarian F, Ajayi C, Hanssen AM, van Sorge NM, Pettersen I, Diep DB, et al. The interaction between Staphylococcus aureus SdrD and desmoglein 1 is important for adhesion to host cells. Sci Rep. 2016;6(1):22134.

58. Pamp SJ, Frees D, Engelmann S, Hecker M, Ingmer H. Spx is a global effector impacting stress tolerance and biofilm formation in Staphylococcus aureus. J Bacteriol. 2006;188(13):4861–70.

59. Singh VK, Syring M, Singh A, Singhal K, Dalecki A, Johansson T. An insight into the significance of the DnaK heat shock system in Staphylococcus aureus. Int J Med Microbiol. 2012;302(6):242–52.

60. Lacey KA, Mulcahy ME, Towell AM, Geoghegan JA, McLoughlin RM. Clumping factor B is an important virulence factor during Staphylococcus aureus skin infection and a promising vaccine target. PLoS Pathog. 2019;15(4).

61. Blau K, Portnoi M, Shagan M, Kaganovich A, Rom S, Kafka D, et al. Flamingo cadherin: a putative host receptor for Streptococcus pneumoniae. J Infect Dis. 2007;195(12):1828–37.

62. Mitchell AM, Mitchell TJ. Streptococcus pneumoniae: virulence factors and variation. Clin Microbiol Infect. 2010;16(5):411–8.

63. Bethe G, Nau R, Wellmer A, Hakenbeck R, Reinert RR, Heinz HP, et al. The cell wall-associated serine protease PrtA: a highly conserved virulence factor of Streptococcus pneumoniae. FEMS Microbiol Lett. 2001;205(1):99–104.

64. Durmort C, Brown JS. Streptococcus pneumoniae lipoproteins and ABC transporters. In: Brown JS, Hammerschmidt S, Orihuela CJ, editors. Streptococcus pneumoniae. London: Elsevier; 2015. p. 181–206.

65. Haas W, Sublett J, Kaushal D, Tuomanen EI. Revising the role of the pneumococcal vex-vncRS locus in vancomycin tolerance. J Bacteriol. 2004;186(24):8463–71.

66. Salter SJ, Cox MJ, Turek EM, Calus ST, Cookson WO, Moffatt MF, et al. Reagent and laboratory contamination can critically impact sequence-based microbiome analyses. BMC Biol. 2014;12:87.

67. Lv L, Wan M, Wang C, Gao X, Yang Q, Partridge SR, et al. Emergence of a plasmid-encoded resistance-nodulation-division efflux pump conferring resistance to multiple drugs, including tigecycline, in Klebsiella pneumoniae. mBio. 2020;11(2).

68. Highlander SK, Hultén KG, Qin X, Jiang H, Yerrapragada S, Mason EO, et al. Subtle genetic changes enhance virulence of methicillin-resistant and -sensitive Staphylococcus aureus. BMC Microbiol. 2007;7(1):99.

69. Gao X, Wang C, Lv L, He X, Cai Z, He W, et al. Emergence of a novel plasmid-mediated tigecycline resistance gene cluster, tmexCD4-toprJ4, in Klebsiella quasipneumoniae and Enterobacter roggenkampii. Microbiol Spectr. 2022;10(4).

70. Shi M, Zhao S, Yu B, Wu WC, Hu Y, Tian JH, et al. Total infectome characterization of respiratory infections in pre-COVID-19 Wuhan, China. PLoS Pathog. 2022;18(2).

71. Graf EH, Simmon KE, Tardif KD, Hymas W, Flygare S, Eilbeck K, et al. Unbiased detection of respiratory viruses by use of RNA sequencing-based metagenomics: a systematic comparison to a commercial PCR panel. J Clin Microbiol. 2016;54(4):1000–7.

72. Rudan I, O’Brien KL, Nair H, Liu L, Theodoratou E, Qazi S, et al. Epidemiology and etiology of childhood pneumonia in 2010: estimates of incidence, severe morbidity, mortality, underlying risk factors and causative pathogens for 192 countries. J Glob Health. 2013;3(1):010401.

73. Rudan I, Boschi-Pinto C, Biloglav Z, Mulholland K, Campbell H. Epidemiology and etiology of childhood pneumonia. Bull World Health Organ. 2008;86:408–16.

74. Khawcharoenporn T, Apisarnthanarak A, Mundy LM. Non-neoformans cryptococcal infections: a systematic review. Infection. 2007;35:51–8.

75. Chen Y, Zhu L, Qian F, Cai H, Yin J. Cryptococcus albidus infected pulmonary mycosis with miliary nodules in CT imaging: two case reports. Respir Med Case Rep. 2024;51:101667.

76. Morgene MF, Botelho-Nevers E, Grattard F, Pillet S, Berthelot P, Pozzetto B, et al. Staphylococcus aureus colonization and non-influenza respiratory viruses: interactions and synergism mechanisms. Virulence. 2018;9(1):1354–63.

77. Dissanayake E, Brockman-Schneider R, Stubbendieck R, Currie C, Gern J. Staphylococcus aureus increases rhinovirus replication and synergistically enhances cytotoxicity during co-infection of the airway epithelium. J Allergy Clin Immunol. 2021;147(2).

78. Dissanayake E, Brockman-Schneider RA, Stubbendieck RM, Helling BA, Zhang Z, Bochkov YA, et al. Rhinovirus increases Moraxella catarrhalis adhesion to the respiratory epithelium. Front Cell Infect Microbiol. 2023;12:1073340.

79. Fazel P, Sedighian H, Behzadi E, Kachuei R, Imani Fooladi AA. Interaction between SARS-CoV-2 and pathogenic bacteria. Curr Microbiol. 2023;80(7):223.

80. Petazzoni G, Bellinzona G, Merla C, Corbella M, Monzillo V, Samuelsen Ø, et al. The COVID-19 pandemic sparked off a large-scale outbreak of carbapenem-resistant Acinetobacter baumannii from the endemic strains at an Italian hospital. Microbiol Spectr. 2023;11(2).

81. Chen F, Lin J, Yang W, Chen J, Qian X, Yan T, et al. Secondary bacterial infections of carbapenem-resistant Acinetobacter baumannii in patients with COVID-19 admitted to Chinese ICUs. BMC Microbiol. 2025;25(1):319.

82. Chotiprasitsakul D, Ao-udomsuk K, Santanirand P. Impact of COVID-19 on epidemiology and mortality risk factors in patients with carbapenem-resistant Acinetobacter baumannii bloodstream infections in a tertiary care hospital in Thailand. J Glob Antimicrob Resist. 2025;43:155–61.

83. Rangel K, Chagas TPG, De-Simone SG. Acinetobacter baumannii infections in times of COVID-19 pandemic. Pathogens. 2021;10(8):1006.

84. Hu L, Han B, Tong Q, Xiao H, Cao D. Detection of eight respiratory bacterial pathogens based on multiplex real-time PCR with fluorescence melting curve analysis. Can J Infect Dis Med Microbiol. 2020;2020:1–11.

85. Jama-Kmiecik A, Frej-Mądrzak M, Sarowska J, Choroszy-Król I. Pathogens causing upper respiratory tract infections in outpatients. Adv Exp Med Biol. 2016;934:89–93.

86. Tang Z, Fan H, Tian Y, Lv Q. Epidemiological characteristics of six common respiratory pathogen infections in children. Microbiol Spectr. 2025;13(7).

87. Jaggi TK, Agarwal R, Tiew PY, Shah A, Lydon EC, Hage CA, et al. Fungal lung disease. Eur Respir J. 2024;64(5):2400803.

88. Li Z, Lu G, Meng G. Pathogenic fungal infection in the lung. Front Immunol. 2019;10:1654.

89. Ojala T, Kankuri E, Kankainen M. Understanding human health through metatranscriptomics. Trends Mol Med. 2023;29(5):376–89.

90. Rajagopala SV, Bakhoum NG, Pakala SB, Shilts MH, Rosas-Salazar C, Mai A, et al. Metatranscriptomics to characterize respiratory virome, microbiome, and host response directly from clinical samples. Cell Rep Methods. 2021;1(6):100091.

91. Nafea AM, Wang Y, Wang D, Salama AM, Aziz MA, Xu S, et al. Application of next-generation sequencing to identify different pathogens. Front Microbiol. 2024;14:117234.

92. Cevik M, Tate M, Lloyd O, Maraolo AE, Schafers J, Ho A. SARS-CoV-2, SARS-CoV, and MERS-CoV viral load dynamics, duration of viral shedding, and infectiousness: a systematic review and meta-analysis. Lancet Microbe. 2021;2(1)–22.

93. Yang S, Rothman RE. PCR-based diagnostics for infectious diseases: uses, limitations, and future applications in acute-care settings. Lancet Infect Dis. 2004;4(6):337–48.

94. Gardy JL, Loman NJ. Towards a genomics-informed, real-time, global pathogen surveillance system. Nat Rev Genet. 2018;19(1):9–20.

95. Struelens MJ, Ludden C, Werner G, Sintchenko V, Jokelainen P, Ip M. Real-time genomic surveillance for enhanced control of infectious diseases and antimicrobial resistance. Front Sci. 2024;2:101.

96. Hasan GM, Mohammad T, Shamsi A, Sohal SS, Hassan MdI. Applications of genome sequencing in infectious diseases: from pathogen identification to precision medicine. Pharmaceuticals. 2025;18(11):1687.

