## Supplementary Figure 1 for "Metatranscriptomics reveals diverse microbial communities in PCR-negative severe acute respiratory infections"

***Rhinovirus A* - 15HB0155 03/2015**

7071 bp

HRV forward primer
CTAGCCTGCGTGGC

5' GCAGATGAGGCTAGAAATTCCCCACTGGCGACAGTGTTCTAGCCTGCGTGGCTGCCTGCGTGTCCTAACGGACACGAAGCCATATACTTGACAAGGTGTG 400
CGTCTACTCCGATCTTTAAGGGGTGACCGCTGTCACAAGATCGGACGCACCGACGGACGCACAGGATTGCCTGTGCTTCGGTATATGAACTGTTCCACAC

Bpu10I

AAGAGCCCCGTGTGCTCACTTTGAGTCCTCCGGCCCCTGAATGTGGCTAACCTTAACCCTGCAGCTAAGGCACACAACCCAGTGTGAACTTAGTCGTAAT 500
TTCTCGGGGCACACGAGTGAAACTCAGGAGGCCGGGGACTTACACCGATTGGAATTGGGACGTCGATTCCGTGTGTTGGGTCACACTTGAATCAGCATTA

GAGCAATTGCGGGATGGGACCAACTACTTTGGGTGTCCGTGTTTCACTTTTTACCTTTTATAATGCTTATGGTGACAATATATAATAATTTGTATATTGG 600
CTCGTTAACGCCCTACCCTGGTTGATGAAACCCACAGGCACAAAGTGAAAAATGGAAAATATTACGAATACCACTGTTATATATTATTAAACATATAACC 3'

ATGAAACCCACAGGCACAAAG
HRV reverse primer

***Rhinovirus B* - 14HB2201 11/2014**

7140 bp

EaeI

HRV forward primer
CTAGCCTGCGTGGC

5' CCCTCCACTAGTTTGGTCGATGAGGCTAGGAATTCCCCACGGGTGACCGTGTCCTAGCCTGCGTGGCGGCCAACCCAGCATATGCTGGGACGCCTTTTTA 400
GGGAGGTGATCAAACCAGCTACTCCGATCCTTAAGGGGTGCCCACTGGCACAGGATCGGACGCACCGCCGGTTGGGTCGTATACGACCCTGCGGAAAAAT

CAGACATGGTGCGAAGACTCGCATGTGCTTGGTTGTGAGTCCTCCGGCCCCTGAATGCGGCTAACCTTAACCCTGGAGCCTTGCGTCACAATCCAGTGAT 500
GTCTGTACCACGCTTCTGAGCGTACACGAACCAACACTCAGGAGGCCGGGGACTTACGCCGATTGGAATTGGGACCTCGGAACGCAGTGTTAGGTCACTA

GGTAAGGTCGTAATGAGCAATTCCGGGATGGGACCGACTACTTTGGGTGTCCGTGTTTCTCATTTTTCTTATTATTGTCTTATGGTCACAGCATATATAT 600
CCATTCCAGCATTACTCGTTAAGGCCCTACCCTGGCTGATGAAACCCACAGGCACAAAGAGTAAAAAGAATAATAACAGAATACCAGTGTCGTATATATA 3'

ATGAAACCCACAGGCACAAAG
HRV reverse primer

***Rhinovirus C* - 15HB0854 7/2015**

7073 bp

HRV forward primer
C TAGCC T GCGTGGT

5' ATCAGGGCAATGGGGCTACACAATCCCCACTGGCGACAGTGGTGTAGCCCGCGTGGTGCCCCGCCCAGGCTTTGCCTGGGACGCCTTAGGAAAGACAGGG 400
TAGTCCCGTTACCCCGATGTGTTAGGGGTGACCGCTGTCACCACATCGGGCGCACCACGGGGCGGGTCCGAAACGGACCCTGCGGAATCCTTTCTGTCCC

FspI

TGTGAAGACCTTAATGCGCTACGTGTGAGTCCTCCGGCCCCTGAATGCGGCTAATCCTAACCCCGTAGCCATGGTGCGCAATCCAGCGCATGTGTGGTCG 500
ACACTTCTGGAATTACGCGATGCACACTCAGGAGGCCGGGGACTTACGCCGATTAGGATTGGGGCATCGGTACCACGCGTTAGGTCGCGTACACACCAGC

TAACGAGTAATTACGGGATGGAACCGACTACTTTGGGTGTCCGTGTTTCACTTTTTATTTGAAATTGTTTCATGGTTACAGATTTGTGTATACAGATTTG 600
ATTGCTCATTAATGCCCTACCTTGGCTGATGAAACCCACAGGCACAAAGTGAAAAATAAACTTTAACAAAGTACCAATGTCTAAACACATATGTCTAAAC 3'

ATGAAACCCACAGGCACAAAG
HRV reverse primer

***Rhinovirus C* - 17HB0238 5/2017**

7062 bp

BglI

HRV forward primer
CCAGCC T GCGTGGC

5' CAGATGGGGCTGGAAATTCCCCACTGGTGACAGTGTTCCAGCCCGCGTGGCTGCCTGTCCCCCTTTCTTGGGGGAAAGCCAATATATAGACAAGGTGCCA 400
GTCTACCCCGACCTTTAAGGGGTGACCACTGTCACAAGGTCGGGCGCACCGACGGACAGGGGGAAAGAACCCCCTTTCGGTTATATATCTGTTCCACGGT

AGAGCCGCGTTTGCTAATTGTGAGTCCTCCGGCCCCTGAATGCGGCTAATCCAACCCCACAGCTATCGCATGGATTCCACCATGTAGGTAGTCGTAATGG 500
TCTCGGCGCAAACGATTAACACTCAGGAGGCCGGGGACTTACGCCGATTAGGTTGGGGTGTCGATAGCGTACCTAAGGTGGTACATCCATCAGCATTACC

GCAACTGTGGGACGGAACCGACTACTTTGGGTGTCCGTGTTTCCTTTTATACCTTATTTTATCTCATGGTGACAACATATATTGATAAGTATATGTTGAC 600
CGTTGACACCCTGCCTTGGCTGATGAAACCCACAGGCACAAAGGAAAATATGGAATAAAATAGAGTACCACTGTTGTATATAACTATTCATATACAACTG 3'

ATGAAACCCACAGGCACAAAG
HRV reverse primer
