## Supplementary figures and images for "Metatranscriptomics reveals diverse microbial communities in PCR-negative severe acute respiratory infections"

### Supplementary Figure 2

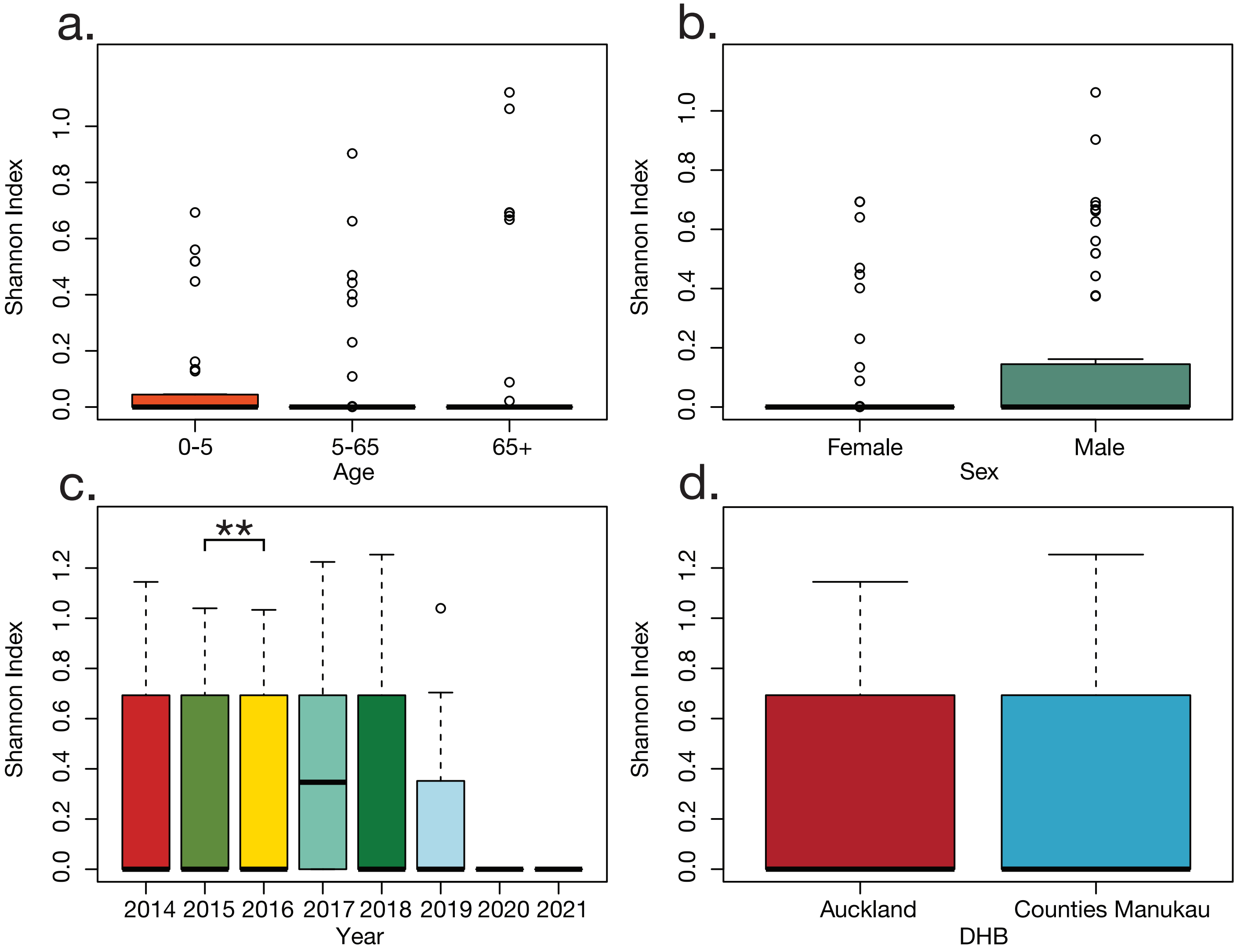
