## Supplementary Table 1 for "Metatranscriptomics reveals diverse microbial communities in PCR-negative severe acute respiratory infections"

**Table S1**. Summary of detected microbial species, number of libraries, and abundance.

| Species | Number of libraries | Total abundance |
| --- | --- | --- |
| Staphylococcus aureus | 20 | 24520.6768 |
| Streptococcus pneumoniae | 34 | 220.806732 |
| Acinetobacter baumannii | 8 | 148.23984 |
| Klebsiella pneumoniae | 2 | 16.0598538 |
| Moraxella catarrhalis | 19 | 3187.42083 |
| Haemophilus influenzae | 4 | 26.3438117 |
| Pseudomonas aeruginosa | 22 | 60.5316385 |
| Serratia marcescens | 1 | 453377.174 |
| Acinetobacter calcoaceticus | 2 | 58.3821887 |
| Rhinovirus C | 10 | 7569.7719 |
| Rhinovirus A | 13 | 11152.1703 |
| Human coronavirus NL63 | 2 | 1800.59207 |
| Parechovirus A | 1 | 20657.4474 |
| Human parainfluenza virus 4b | 1 | 11863.1738 |
| Rhinovirus B | 1 | 471.239369 |
| Human papillomavirus | 2 | 22.8309337 |
| Human coronavirus OC43 | 6 | 369531.961 |
| Human coronavirus HKU1 | 2 | 13464.3308 |
| Human coronavirus 229E | 1 | 8.74330824 |
| Human alphaherpesvirus 3 | 1 | 213.968102 |
| Gammaherpesvirus 4 | 1 | 6.01903989 |
| Measles virus genotype B3 | 1 | 591.50171 |
| Purpureocillium lilacinum | 2 | 54961.596 |
| Penicillium chrysogenum | 7 | 29169.3165 |
| Acremonium sclerotigenum | 2 | 28037.1343 |
| Naganishia albida | 9 | 170643.804 |
